# The NLRP3 inflammasome selectively drives IL-1β secretion by *Pseudomonas aeruginosa* infected neutrophils and regulates bacterial killing *in vivo*

**DOI:** 10.1101/2022.04.07.487503

**Authors:** Martin S. Minns, Karl Liboro, Tatiane S. Lima, Serena Abbondante, Brandon A. Miller, Michaela E. Marshall, Jolynn Tran-Chau, Arne Rietsch, George R. Dubyak, Eric Pearlman

**Author notes:** T.L. current address: Department of Biological Sciences, California State Polytechnic University, Pomona, CA. These authors contributed equally (Martin S. Minns, Karl Liboro, Tatiane S. Lima).

## Abstract

Macrophages infected with Gram-negative bacteria expressing Type III secretion system (T3SS) activate the NLRC4 inflammasome, resulting in Gasdermin D (GSDMD)-mediated IL-1β secretion and pyroptosis. Here we examined inflammasome signaling in neutrophils infected with *Pseudomonas aeruginosa* strain PAO1 that expresses the T3SS effectors ExoS and ExoT. IL-1β secretion by neutrophils required the T3SS needle and translocon proteins and GSDMD. In macrophages, PAO1 and mutants lacking ExoS and ExoT (*ΔexoST*) stimulated NLRC4 for IL-1β secretion. While IL-1β release from *ΔexoST* infected neutrophils was also NLRC4-dependent, this was redirected to NLRP3-dependence by PAO1 infection via the ADP ribosyl transferase activity of ExoS. Genetic and pharmacologic approaches revealed that NLRP3, but not NLRC4, was essential for bacterial killing and limiting disease severity in a murine model of *P. aeruginosa* corneal infection. This reveals a novel role for ExoS ADPRT in regulating inflammasome subtype usage by neutrophils versus macrophages and an unexpected role for NLRP3 in *P. aeruginosa* keratitis.

---

The discovery of the Gasdermin (GSDM) family of pore forming proteins has greatly increased our understanding of the mechanism of cell death in inflammation, cancer, and autoimmunity in addition to host defense, and has identified potential new targets for pharmacological intervention ^1, 2, 3^. Gasdermin D (GSDMD) is a critical mediator of inflammasome-induced release of bioactive IL-1β and pyroptosis in myeloid leukocytes. Compared with macrophages, there are relatively few studies examining the role of GSDMD or other Gasdermin family members in neutrophils, even though neutrophils vastly outnumber other cells recruited to sites of inflammation and infection, especially at early time points. We and others reported that neutrophils are a major source of IL-1β in response to either canonical NLRP3 activators such as ATP, by *Streptococcus pneumoniae* activation of NLRP3, or through NLRC4 driven by *Salmonella* infections, and that in contrast to macrophages, IL-1β secretion by neutrophils occurs in the absence of GSDMD mediated pyroptotic cell death ^4, 5, 6, 7, 8, 9^.

*P. aeruginosa* is a ubiquitous environmental Gram-negative bacterium, and is an important cause of dermal, ocular, and pulmonary disease, including cystic fibrosis. The Type III secretion system (T3SS) is a major virulence factor of *P. aeruginosa* that includes a complex needle structure that penetrates the plasma membrane of the target mammalian cell and injects effector proteins directly into the cytosol. Up to four effectors (exoenzymes) with distinct functions are injected via the T3SS, including cell lysis by the phospholipase ExoU, modulation of protein function by ADP ribosyl transferase (ADPRT) or GTPase activating protein (GAP) activities of ExoS and ExoT ^10, 11^.

Our studies have focused on blinding *P. aeruginosa* infections of the cornea (keratitis), which are a major cause of corneal ulcers in the USA and worldwide ^12, 13^, and most clinical isolates from *P. aeruginosa* infected corneas express ExoS and ExoT rather than ExoU ^14, 15^. Quantitative PCR analysis of neutrophil rich corneal ulcers caused by *P. aeruginosa* or *S. pneumoniae* revealed elevated gene expression of IL-1β, NLRC4, NLRP3 and ASC compared with uninfected human corneas ^15^.

Using a murine model of *P. aeruginosa* keratitis, we reported that an intact T3SS and the ADPRT activities of ExoS and ExoT for virulence in infected corneas ^16^. We also showed that ExoS ADP-ribosylates the RAS small GTPase in neutrophils and blocks assembly of the NADPH oxidase complex, resulting in lower ROS production and impaired bacterial killing ^17^.

Both pyroptosis and IL-1β secretion in macrophages are coordinately driven by GSDMD, which is cleaved by caspase-1 (or caspase-11) to generate N-terminal subunits (N-GSDMD), which in turn form a pore in the plasma membrane leading to the release of cleaved IL-1β^15, 18^. We found that instead of inserting into the neutrophil plasma membrane, N-GSDMD induced by the NLRP3 inflammasome localizes to primary granules and autophagosomes, leading to release of neutrophil elastase into the cytosol and cleavage of pro-GSDMD by this protease ^8^.

Distinct roles for Gasdermin family proteins in neutrophils versus macrophages also applies to GSDME, which was first reported as a mutated gene (DFNA5) associated with hearing loss, but also forms plasma membrane pores following cleavage by caspase-3 ^1^. The *Yersinia* T3SS acetyltransferase *YopJ* activates GSDME rather than GSDMD via a RIPK1/caspase-3 pathway to induce IL-1β secretion and cell death in neutrophils ^19^.

In the current study, we characterized the relative contributions of T3SS, the NLRP3, and NLRC4 inflammasomes, and GSDMD and GSDME to IL-1β release and pyroptosis in neutrophils versus macrophages following infection with *P. aeruginosa*. Surprisingly, we found that the ExoS ADP ribosyltransferase activity led to IL-1β secretion through the NLRP3 rather than the NLRC4 inflammasome in neutrophils, but not macrophages. Consistent with these in vitro findings, NLRP3 and caspase-1 were required to regulate *P. aeruginosa* growth and disease severity in infected corneas, but there was no apparent role for the NLRC4 inflammasome.

Finally, we show that although GSDMD is reported to mediate formation of neutrophil extracellular traps (NETosis) in response to PMA or intracellular LPS ^20, 21^, we found that NET accumulation induced by PMA or *P. aeruginosa* was completely independent of GSDMD. Overall, these observations underscore how inflammasome signaling networks are differentially engaged by the *P. aeruginosa* T3SS in neutrophils versus macrophages and illustrate their relative contributions to disease pathogenesis.

## RESULTS

### The P. aeruginosa T3SS needle and translocon structures mediate neutrophil pyroptotic IL-1β release through GSDMD

**Figure 1A** shows the major proteins that comprise the *P. aeruginosa* T3SS, including the injectosome (the needle structure) and the translocon pore that penetrates the host cell membrane. These transport the effector exoenzymes from the bacterial cytosol through the inner and outer membranes and into the target eukaryotic cell. To examine the role of T3SS in IL-1β secretion, bone marrow neutrophils from C57BL/6 mice were isolated by negative bead selection (>95% purity), incubated 3h with LPS to induce pro-IL-1β, and infected (1h at an MOI of 30:1) with *P. aeruginosa* strain PAO1 that expresses ExoS, T and Y, with Δ*pscD* (apparatus mutant), Δ*popB* (translocon mutant), or with exoenzyme mutants Δ*exoSTY* and Δ*exoST* that express the needle structure, but not the exoenzymes.

**Figure 1.**
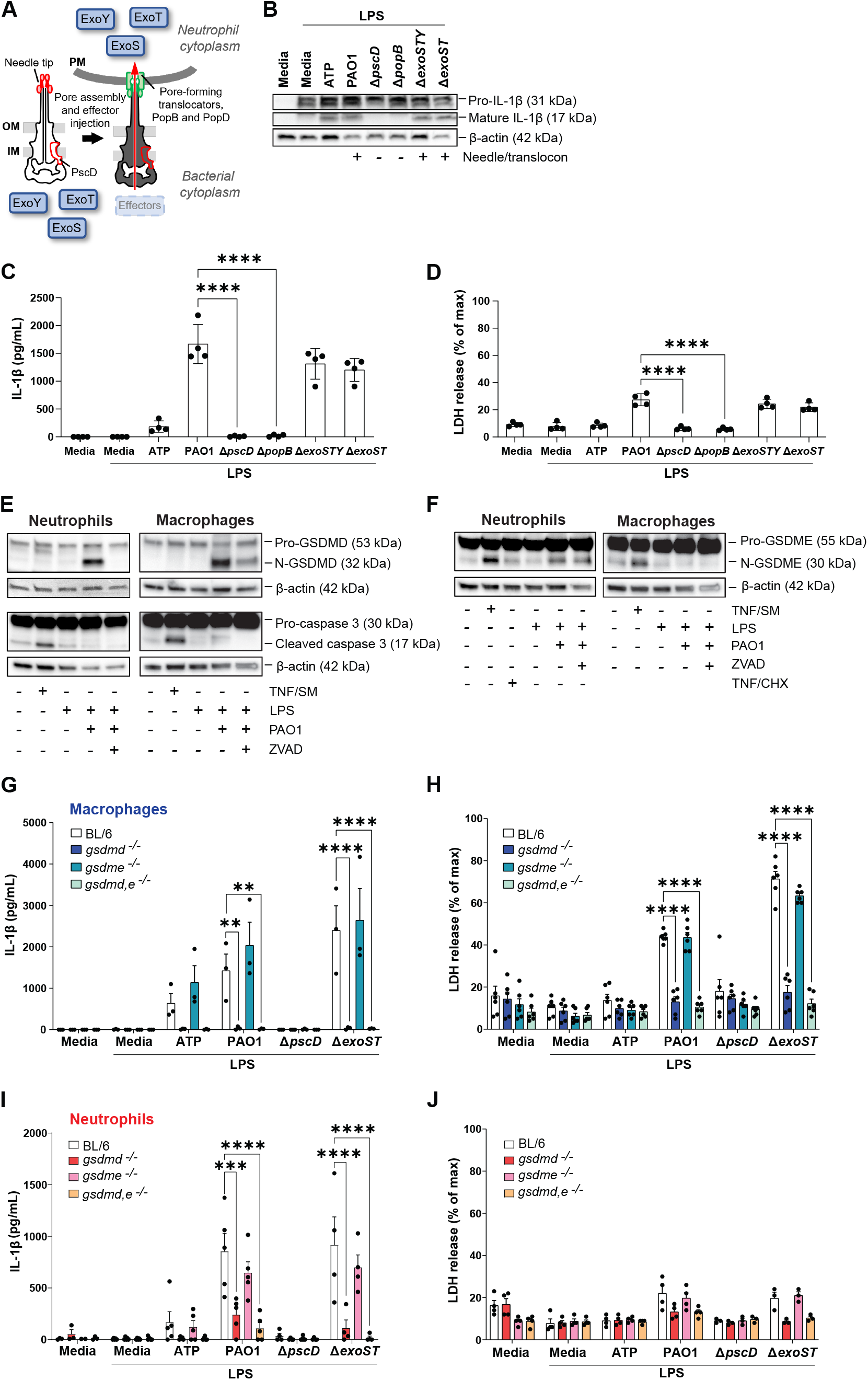
The role of T3SS, GSDMD and GSDME in IL-1β secretion by *P. aeruginosa* infected neutrophils. ***A*.** *P. aeruginosa* T3SS is preassembled and spans the bacterial cell envelope (IM, inner membrane, OM, outer membrane and PG, peptidoglycan layer). Following cell contact, the pore-forming translocator proteins, PopB (B) and PopD (D), assemble into a pore, which docks to the needle tip (PcrV, red). Effector secretion is triggered, resulting in export of effector proteins across the bacterial cell envelope and host cell plasma membrane (PM) into the cytosol of the targeted cell. *P. aeruginosa* strain PAO1 used in the current study expresses ExoS, ExoT, and ExoY, but not ExoU. **B.** GSDMD and IL-1β cleavage in lysates or combined supernatants and lysates from bone marrow neutrophils from C57BL/6 mice primed 3h with LPS and incubated 45 min with ATP or with *P. aeruginosa* strain PAO1 or PAO1 mutants Δ*pscD* (no needle), Δ*popB* (no translocon), Δ*ExoSTY* or Δ*ExoST*. **C-E:** Bone marrow neutrophils from C57BL/6 or GSDMD^-/-^ mice incubated with PAO1 or the same mutants. Quantification of IL-1β(**C**), LDH (**D**) and CXCL2 (**E**). Each data point is the mean of independent biological repeat experiments. Statistical significance was assessed by 1-way ANOVA followed by Tukey posttest. Western blots are representative of 3 repeat experiments.

Pro-IL-1β was cleaved to the bioactive 17kD IL-1β in neutrophils infected with *P. aeruginosa* strains PAO1, Δ*exoSTY* and Δ*exoST*, but not in neutrophils infected with the Δ*pscD* apparatus mutant or the Δ*popB* translocon mutant (**Figure 1B).** Consistent with this finding, only strains expressing the intact needle structure (PAO1, Δ*exoSTY* and Δ*exoST*) were found to induce IL-1β secretion (**Figure 1C).** LDH release, although higher in PAO1 infected neutrophils was <30% of the lysis buffer control (**Figure 1D)**. Flagellin also induces IL-1β secretion by macrophages ^22, 23, 24^; however, there were no significant differences between PAO1 and a Δ*fliC* mutant in the activation of IL-1β secretion or LDH release by neutrophils (**Figure S1A, B)**. Notably, casein-elicited peritoneal neutrophils that were infected for 1h with PAO1 or *P. aeruginosa* mutant strains in the absence of LPS priming released IL-1β with similar strain-dependent differences as was observed in LPS-primed bone marrow neutrophils (**Figure S1C,D**).

Canonical ATP - induced activation of NLRP3 mediated IL-1β secretion in neutrophils is dependent on GSDMD, which in contrast to macrophages, does not lead to accumulation of plasma membrane N-GSDMD pores and pyroptotic cell death ^8^. To determine the role of GSDMD and GSDME in *P. aeruginosa* – induced IL-1β secretion, bone marrow neutrophils and bone marrow derived macrophages from C57BL/6, *Gsdmd^-/-^*, *Gsdme^-/-^* and *Gsdmd^-/-^/Gsdme^-/-^* mice were primed for 3h with LPS and infected with PAO1 strain for 1h prior to analysis of pro-GSDMD and pro-GSDME cleavage. As a positive control for caspase-3 and pro-GSDME cleavage, neutrophils and macrophages were stimulated with TNF-α plus a second mitochondria-derived activator of caspases (SMAC) mimetic (SM), which activate the ripoptosome and apoptotic caspases that mediate GSDME cleavage ^25^ or with TNF+ cycloheximide (CHX) as described ^19^. Cells were also stimulated in the presence of the pancaspase inhibitor ZVAD, and macrophages were incubated in the presence of glycine to inhibit cell lysis. GM-CSF was added to neutrophil cultures to prevent spontaneous caspase-3 mediated apoptosis.

The 31 kDa N-GSDMD cleavage product accumulated in PAO1 infected neutrophils and macrophages via a ZVAD-inhibited process, whereas caspase-3 and caspase-8 cleavage was only detected following TNF/SM stimulation of the ripoptosome **(Figure 1E, S1E)**. GSDME cleavage was induced by TNF/SM in neutrophils and macrophages in a similar manner as caspase-3, whereas there was no stimulatory effect of TNF/CHX; however, PAO1 infection resulted in GSDME processing in neutrophils, but not in macrophages (**Figure 1F**). Notably, the PAO1-induced accumulation of cleaved GSDME in neutrophils was not inhibited by ZVAD. Taken together with the absence of caspase-3 processing by PAO1, these findings indicate that PAO1 infected neutrophils cleave pro-GSDME by a caspase-independent mechanism.

To ascertain if GSDMD and GSDME are required for IL-1β secretion following infection with ExoS expressing *P. aeruginosa*, bone marrow neutrophils isolated from *Gsdmd^-/-^*, *Gsdme^-/-^* and *Gsdmd^-/-^*/*Gsdme^-/-^* mice were primed with LPS and incubated 1h with ATP, PAO1, Δ*pscD* or Δ*exoST* mutant bacteria. We found that IL-1β secretion and LDH release by macrophages stimulated with ATP, PAO1 or Δ*exoST* mutants were significantly lower in *Gsdmd^-/-^* and *Gsdmd^-/-^*/*Gsdme^-/-^* macrophages compared with C57BL/6 neutrophils; however, there was no significant difference in IL-1β secretion between macrophages from C57BL/6 and *Gsdme^-/-^* mice **(Figure 1G, H)**. Similarly, IL-1β secretion and LDH release by neutrophils simulated with ATP or infected with PAO1 or Δ*exoST* was completely dependent on GSDMD with no obvious contribution from GSDME **(Figure 1I, J)**.

Collectively, these findings demonstrate that *P. aeruginosa* induces IL-1β secretion and low levels of cell death in neutrophils, both of which are dependent on expression of the needle structure and the translocon. As Δ*exoST* mutants induced similar responses to wild type PAO1, we initially conclude that T3SS exotoxins are not required for IL-1β release. Further, although GSDMD and GSDME are processed in PAO1 infected neutrophils and macrophages, and GSDMD is required for IL-1β secretion and LDH release by both myeloid cell types, there is no apparent role for GSDME in IL-1β secretion induced by T3SS expressing *P. aeruginosa*.

### Neutrophil IL-1β secretion is NLRC4 independent following infection with ExoS/ExoT expressing *P. aeruginosa*

In macrophages, IL-1β secretion and pyroptosis induced by the needle structure and flagella of *P. aeruginosa* and other Gram-negative bacteria is mediated by NLRC4 ^23, 26, 27^. As the *P. aeruginosa* needle structure is required for GSDMD cleavage and IL-1β secretion by neutrophils, we next examined if this is dependent on NLRC4. Bone marrow neutrophils and bone marrow derived macrophages from C57BL/6 or *Nlrc4^-/-^* mice were LPS primed and infected 1h with PAO1, Δ*pscD* or *ΔexoST*, and IL-1β and LDH were quantified as before.

IL-1β secretion and LDH release were elevated in C57BL/6 and *Nlrc4^-/-^* macrophages stimulated with the canonical NLRP3 activator ATP; however, PAO1 and Δ*exoST* induced elevated IL-1β and LDH in C57BL/6, but not *Nlrc4^-/-^* macrophages (**Figure 2A, B)**. Similarly, IL-1β secretion and LDH release in neutrophils infected with Δ*exoST* mutants was NLRC4 dependent; however in contrast to macrophages, *Nlrc4^-/-^* neutrophils infected with PAO1 secreted IL-1β and released LDH in amounts not significantly different from C57BL/6 neutrophils (**Figure 2C,D**).

**Figure 2.**
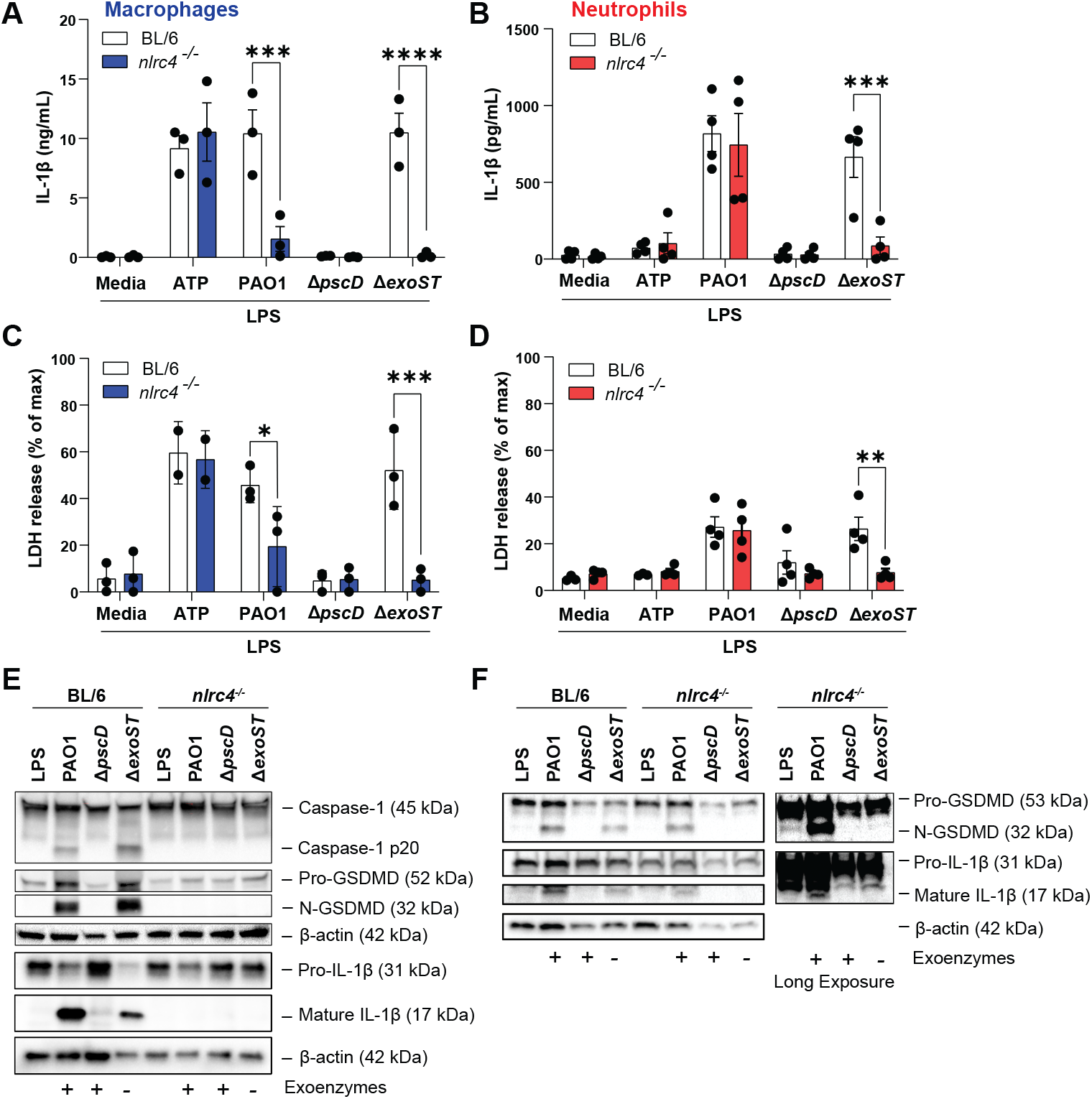
PAO1 mediated IL-1β secretion by neutrophils is NLRC4 independent. Bone marrow derived macrophages (**A-C)** and bone marrow neutrophils (**D-F)** from C57BL/6 and NLRC4^-/-^ mice were incubated with PAO1, Δ*pscD* (no needle), Δ*popB* (no translocon), or Δ*ExoST*. IL-1β secretion and LDH release were quantified (**A,B, D, E**) and GSDMD and IL-1b cleavage were examined by western blot (**C, F**). Each data point is the mean of independent biological repeat experiments. Statistical significance was assessed by 1-way ANOVA followed by Tukey post-test. Western blots are representative of 3 repeat experiments.

In support of these findings, NLRC4 was required for IL-1β, caspase-1 and GSDMD cleavage in macrophages (**Figure 2E)**; however, while was no difference in IL-1β and GSDMD cleavage in PAO1-infected *Nlrc4^-/-^* compared with C57Bl/6 neutrophils, NLRC4 was required for IL-1β and GSDMD cleavage when neutrophils were infected with the Δ*exoST* mutant (**Figure 2F).**

Collectively, these findings are aligned with earlier reports that in macrophages, *P. aeruginosa* induced IL-1β secretion is dependent on NLRC4 inflammasome signaling ^26, 27^; however, we now show that in neutrophils infected with *P. aeruginosa* expressing ExoS and ExoT (PAO1), GSDMD cleavage and IL-1β secretion is NLRC4 independent.

### Neutrophil IL-1β secretion is NLRP3 dependent in the presence of ExoS ADPRT

As IL-1β is secreted by PAO1 infected neutrophils in the absence of NLRC4, we asked if ExoS and/or ExoT induce IL-1β secretion through the NLRP3 inflammasome. C57BL/6 and *Nlrp3^-/-^* neutrophils and macrophages were infected with either PAO1 or the *ΔexoST* mutant strain, IL-1β secretion and LDH release, as well as the cleavage of pro-IL-1β and GSDMD were assayed.

Consistent with the dominant role for NLRC4 in IL-1β secretion and LDH release in macrophages infected with PAO1 or the *ΔexoST* mutant, there was no difference between *Nlrp3^-/-^* and C57BL/6 macrophages in IL-1β secretion or LDH release, although LPS/ATP induced IL-1β secretion was NLRP3 - dependent (**Figure 3A,B)**.

**Figure 3.**
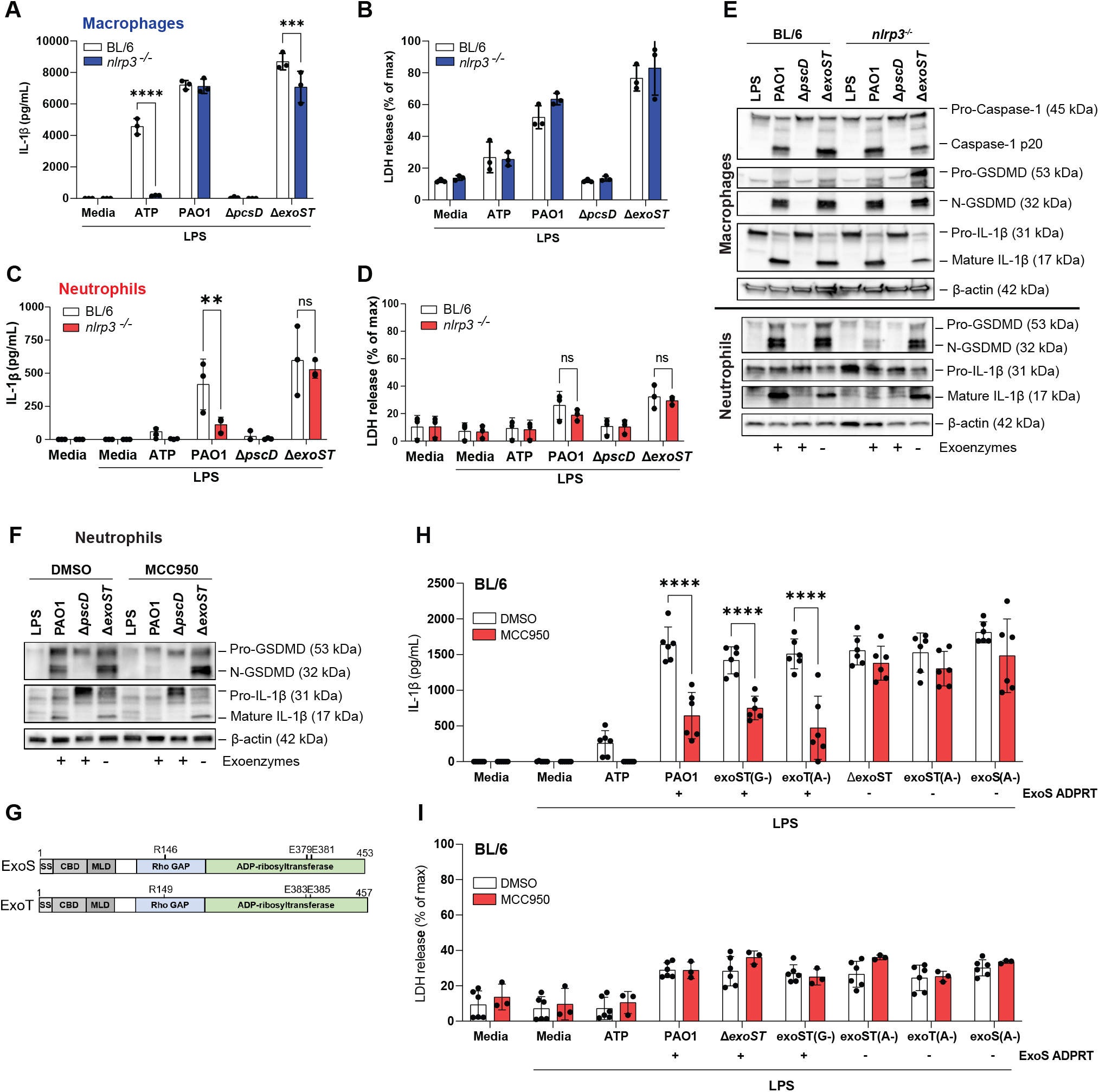
PAO1 mediated IL-1β secretion by neutrophils is NLRP3 dependent in the presence of ExoS ADPRT. Bone marrow derived macrophages (**A,B)** and bone marrow neutrophils (**C,D)** from C57BL/6 and *Nlrp3^-/-^* mice were incubated with PAO1, Δ*pscD* or Δ*exoST*. IL-1β secretion and LDH release were quantified. **E.** GSDMD and IL-1β cleavage were examined by western blot in macrophages and neutrophils from *Nlrp3^-/-^* mice. **F.** GSDMD and IL-1β cleavage in C57BL/6 neutrophils infected with PAO1 or mutants in the presence of the NLRP3 inhibitor MCC950. **G.** GTPase (GAP) and ADP ribosyltransferase (ADPRT) regions of ExoS and ExoT showing amino acids required for enzymatic function and which are also the sites of point mutations. **H.** Neutrophils from C57BL/6 mice infected with PAO1, Δ*exoST* or mutants with indicated point mutations (A-, G-) in the presence of MCC950. Strains expressing ExoS ADPRT are indicated. Each data point is the mean of independent biological repeat experiments. Western blots are representative of 3 repeat experiments. Statistical significance was assessed by 1-way ANOVA followed by Tukey post-test‥

In marked contrast, IL-1β secretion by *Nlrp3^/-^* neutrophils infected with PAO1 was significantly lower compared with C57BL/6 neutrophils, whereas there was no difference between C57BL/6 and *Nlrp3^/-^* neutrophils incubated with the Δ*exoST* mutant **(Figure 3C)**. As in prior experiments, the modest LDH release from infected neutrophils was <30% of the detergent control **(Figure 3D)**. Consistent with the ELISA data, there was no difference in IL-1β, caspase-1 or GSDMD cleavage between C57BL/6 and *Nlrp3^-/-^* macrophages infected with PAO1 or Δ*exoST* **(Figure 3E, upper panel**), whereas IL-1β and GSDMD cleavage was lower in *Nlrp3^-/-^* neutrophils infected with PAO1, but not the ΔexoST mutant (**Figure 3E, lower panel).**

As a second approach to examine the role of NLRP3, neutrophils from C57BL/6 mice were incubated with the well-characterized, highly specific NLRP3 inhibitor MCC950 ^27^ prior to infection with *P. aeruginosa*. As with *Nlrp3^-/-^* neutrophils, GSDMD and IL-1β cleavage was inhibited in MCC950 treated neutrophils following infection with PAO1, whereas there was no effect of MCC950 in neutrophils infected with the Δ*exoST* mutant (**Figure 3F)**. *Nlrc4^-/-^* neutrophils incubated with MCC950 responded the same as C57BL/6 neutrophils, i.e., MCC950 inhibited IL-1β following infection with PAO1 or ExoS ADPRT expressing mutants (**Figure S2A,B**). Collectively, these findings indicate that ExoS and/or ExoT either drive NLRP3 inflammasome usage or inhibit NLRC4 activity in neutrophils, but not in macrophages.

*P. aeruginosa* ExoS and ExoT are bifunctional exoenzymes as shown in **Figure 3G** – a GTPase Activating Protein (GAP) domain that targets small GTPases (Rho, Rac, CDC42) and disrupts host cell cytoskeletal activities, and ADP ribosyltransferase (ADPRT) that post-translationally modifies the activity of multiple proteins ^10^. To identify which of these functions of ExoS or ExoT mediate NLRP3 dependent IL-1β secretion, bone marrow neutrophils from C57BL/6 mice were infected with PAO1 expressing point mutations in the enzymatic regions of either the GAP or ADPRT domains of ExoS or ExoT (indicated in the diagram) in the presence or absence of MCC950.

We found that point mutations in GAP domains of ExoS and ExoT (*exoST(G-)*) or in the ExoT ADPRT domain (*exoT(A-)*) phenocopied PAO1 in showing MCC950 inhibition (NLRP3 activity), whereas point mutations inactivating both ExoS and ExoT ADPRT domains (*exoST(A-*)), or only the ExoS ADPRT domain (*exoS(A-)*) phenocopied the Δ*exoST* mutant strain in showing no effect of MCC950 on readouts of inflammasome activity (**Figure 3H,I)**.

Taken together, these findings identify a requirement for the ADP ribosyl transferase activity of ExoS to redirect inflammasome signaling to NLRP3-dependent rather than NLRC4-dependent pathways during *P. aeruginosa* infection of neutrophils.

In contrast to our results using *Nlrc4^-/-^* neutrophils, a recent study showed that IL-1β secretion and LDH release following a 3h infection with PAO1 (in the absence of LPS priming) was completely dependent on NLRC4 ^28^. Using this protocol, we also found that IL-1β secretion by PAO1 is NLRC4-dependent compared with LPS primed neutrophils (**Figure S2C-F**). As we reported that NLRP3 is transcribed *de novo* in neutrophils in response to NFkB-activating mediators ^6^; therefore, it is likely that in the absence of priming by LPS a 3h PAO1 infection at a 10:1 multiplicity of infection does not induce sufficient NLRP3 expression in bone marrow neutrophils, leaving NLRC4 as the default inflammasome responding to *P. aeruginosa*. The primary role for NLRP3 is supported by our findings that IL-1β secretion by peritoneal neutrophils in the absence of LPS priming NLRP3 dependent (**Figure S2G,H**). The latter model can be considered more physiological because blood neutrophils recruited into the inflammatory peritoneal environment are already primed and are expressing NLRP3.

### *P. aeruginosa* corneal infection (keratitis) is regulated by NLRP3 and Caspase-1

As noted earlier, *P. aeruginosa* is an important cause of painful corneal infections worldwide, resulting in visual impairment and blindness. To characterize the relative contributions of NLRP3 and NLRC4 *in vivo*, we used a well-established murine model of *P. aeruginosa* keratitis in which the corneal epithelium is abraded, and 1×10^4^ GFP expressing bacteria (GFP-PAO1) are added topically in 2 μl saline ^29^. That study also showed that in PAO1 infected C57BL/6 mice, there is early central corneal opacification that is associated with neutrophil infiltration and control of bacterial growth; however, in the absence of IL-1β, bacteria replicate in the central cornea (at the site of infection), leading to perforation within 48-72h postinfection ^29^.

To determine the relative contribution of NLRP3 and NLRC4 inflammasomes in *P. aeruginosa* keratitis, corneas of *Nlrp3^-/-^* and *Nlrc4^-/-^* mice, and C57BL/6 mice given systemic MCC950 were abraded and infected with 1β 10^4^ PAO1, and corneal opacification, GFP intensity and CFU were examined after 24h.

In infected C57BL/6 and *Nlrc4^-/-^* mice, there was central corneal opacity and low levels of GFP; however, when these mice were given systemic MCC950, there was decreased central corneal opacification and increased GFP-bacteria and CFU (**Figure 4A-C)**. *Nlrp3^-/-^* mice had the same phenotype as C57BL/6 and *Nlrc4^-/-^* mice given MCC950. Consistent with a requirement for the NLRP3 inflammasome to regulate bacterial growth, *caspase-1/11^-/-^* and *caspase-1^-/-^* mice had higher GFP intensity and elevated CFU compared with C57BL/6 mice (**Figure 4D-F)**. Infected corneas from MCC950 treated C57BL/6 mice and *Nlrp3^-/-^* mice had significantly less bioactive IL-1β than untreated C57BL/6 corneas, although there was no difference in the number of neutrophils or monocytes in infected corneas of C57BL/6 compared with *Nlrp3^-/-^* mice (**Figure S3A-D**).

**Figure 4.**
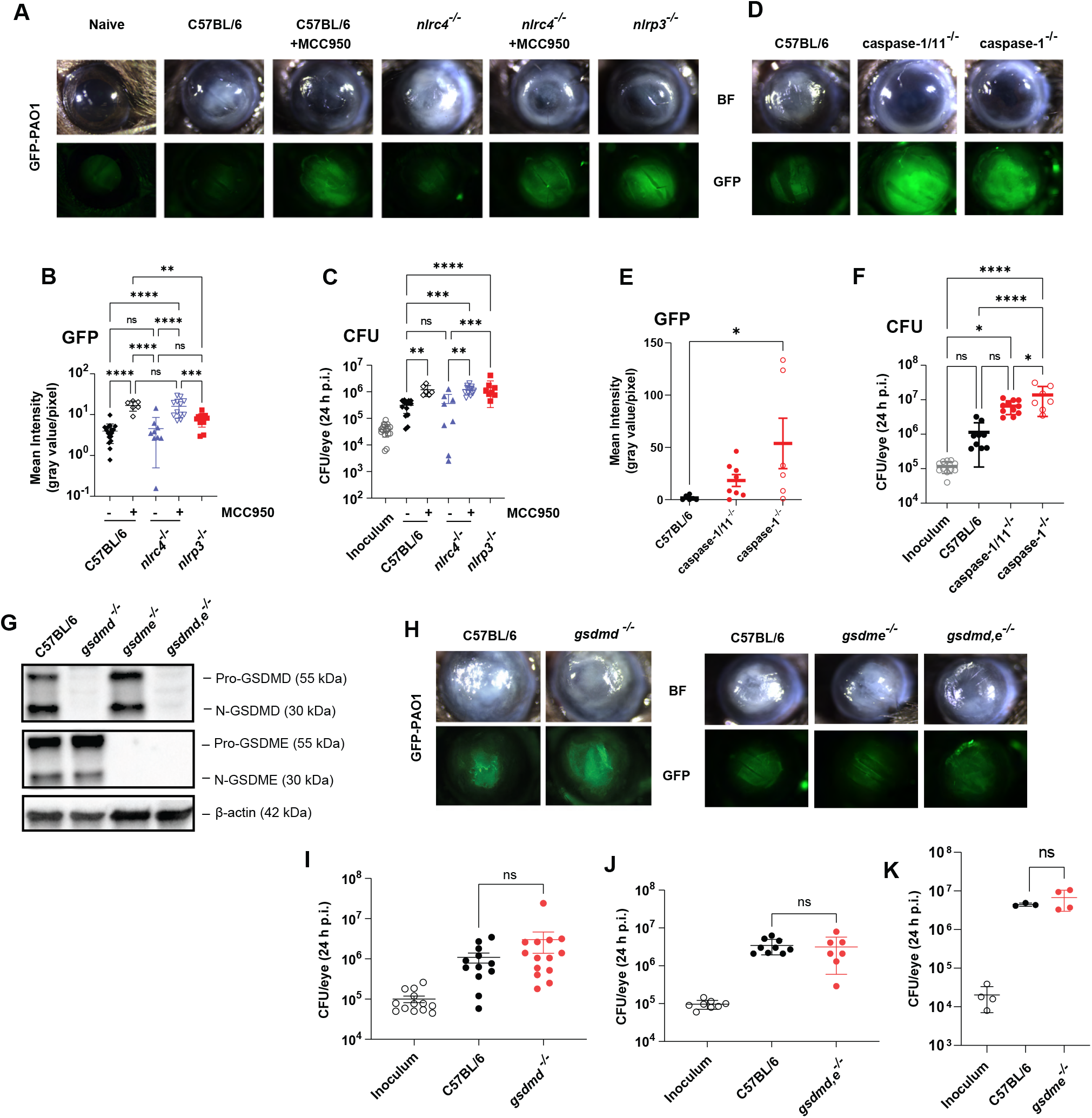
The role of inflammasomes, GSDMD and GSDME in *P. aeruginosa* corneal infections. **A-C.** Corneas of MCC950 treated C57BL/6 and *Nlrc4^-/-^* mice, and *Nlrp3^-/-^* mice were infected with 1×10^5^ GFP expressing PAO1. After 24h, corneal opacification and total GFP bacteria were quantified by image analysis, and viable bacteria were measured by CFU. **A.** representative images of infected corneas and GFP-PAO1. **B.** Quantification of GFP in infected corneas. **C.** CFU in infected corneas. **D-F.** Corneal opacification, GFP-PAO1 and CFU in *caspase-1 and caspase-1/11^-/-^* mice. **G-L.** PAO1 infected C57BL/6, *gsdmd^-/-^, gsdme^-/-^* and *gsdmd^-/-^*/*gsdme^-/-^* corneas. **G.** GSDMD and GSDME cleavage in infected corneas. **H, I.** Representative corneas, and CFU (**J-L**). GSDMD and GSDME cleavage products were examined by detected blot. Each data point is the mean of independent biological repeat experiments. Statistical significance was assessed by 1-way ANOVA followed by Tukey post-test. Western blots are representative of 3 repeat experiments.

Corneal disease and bacterial killing were also assessed in infected *Gsdmd^-/-^*, *Gsdme^-/-^* and *Gsdmd^-/-^*/*Gsdme^-/-^* mice. We found that GSDMD and GSDME were cleaved in infected corneas (**Figure 4G)**; however, there were no significant differences in CFU or corneal disease between infected C57BL/6 and *Gsdmd^-/-^*, *Gsdme^-/-^* or *Gsdmd^-/-^*/*Gsdme^-/-^* mice (**Figure 4H-K)**. There was also no difference in IL-1β secretion or the number of neutrophils and monocytes in infected corneas of C57BL/6 and *Gsdmd^-/-^* mice (**Figure S3E-G**). Finally, we found no role for neutrophil elastase in this process as infected *Elane^-/-^* corneas showed no difference opacification, CFU or IL-1β cleavage from C57BL/6 mice (**Figure S3I-K)**.

Overall, these findings support a role for NLRP3 and caspase-1 in regulating bacterial killing and corneal disease severity with no apparent role for NLRC4 and no requirement for GSDMD, GSDME, or neutrophil elastase.

### Induction of neutrophil extracellular traps (NETs) by P. aeruginosa infection is not dependent on GSDMD or GSDME

Although *P. aeruginosa* induced neutrophils exhibit <30% LDH release following infection, neutrophils also undergo a mode of cell death (NETosis) coordinated with the programmed release of nuclear DNA, histones and cytosolic proteins as neutrophil extracellular traps (NETs) in response to protein kinase C (PKC) activation of neutrophils by phorbol myristate acetate (PMA) or after incubation with bacteria or fungi ^30, 31^. NET proteins include histones, neutrophil proteases, and peptides that have antimicrobial and cytotoxic activities. GSDMD was reported to mediate NETosis induced by PMA ^21^, or infection with intracellular *Salmonella, Citrobacter* or transfected LPS that accumulates in the cytosol with consequent activation of non-canonical caspase-11 (caspases 4 and 5 in human cells) ^20^. Therefore, as *P. aeruginosa* induces NET accumulation in murine models of pulmonary and corneal infections ^30, 31^, we examined if there is a role for GSDMD or GSDME in *ex vivo* NETosis induced by *P. aeruginosa* using either neutrophils from C57BL/6 mice incubated with reported GSDMD inhibitors or using neutrophils from GSDMD^-/-^ and GSDMD^-/-^ mice.

Neutrophils isolated from the peritoneal cavity of C57BL/6 mice following sterile inflammation were incubated with PAO1 at a ratio of 30:1 in the presence of LDC7559, which was reported to inhibit GSDMD-dependent NETosis ^17^, or with two other known GSDMD inhibitors, necrosulfonamide (NSA) and disulfiram ^32, 33^. We found a significant reduction in IL-1β secretion by neutrophils incubated with NSA or disulfiram, but not LDC7559; however, there was no effect on NET formation (as quantified by accumulation of extracellular Sytox Green-DNA fluorescent complexes) by any of these compounds (**Figure 5A,B)**.

**Figure 5.**
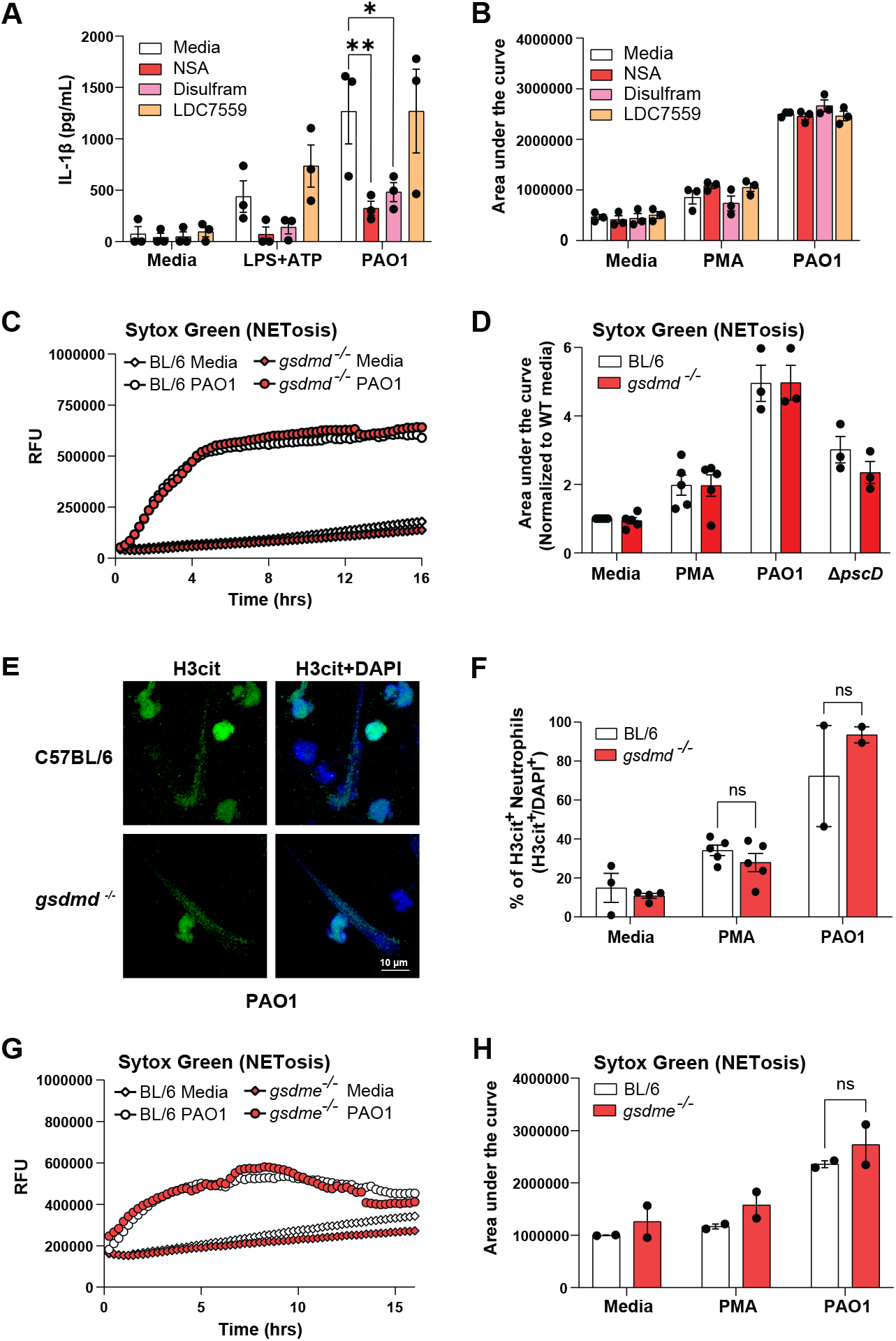
PAO1 induced NETosis is independent of ROS, GSDMD and GSDME. **A, B:** IL-1β secretion (**A**) and NETosis (**B**) by peritoneal neutrophils incubated with reported GSDMD inhibitors necrosulfonamide (NSA), disulfiram, and LDC7559. NETosis was quantified by SYTOX detection of extracellular DNA after 16h (Area under the curve; a representative time course is shown in Figure S4A). **C,D.** Peritoneal neutrophils from C57BL/6 and *Gsdmd^-/-^* mice were incubated 16h with PMA, PAO1 or Δ*pscD* and extracellular DNA was measured by Sytox. **C.** representative time course, and **D.** quantification of the area under the curve. **E.** representative images of citrullinated histone 3 (H3Cit) as an indicator of NETosis in PAO1 infected neutrophils (original and other images are in Figure S4F). **F.** Quantification of H3Cit+ neutrophils per high power field. G, H. Representative time course and quantification of NETosis in C57BL/6 and *Gsdme^-/-^* mice.

As a second approach to examine the role of GSDMD in *P. aeruginosa* - mediated NETosis, peritoneal neutrophils were isolated from C57BL/6 and *Gsdmd^-/-^* mice and infected with PAO1, and NETosis induced by *P. aeruginosa* was quantified using either Sytox Green, or antibodies to citrullinated histone 3 (H3Cit). H3Cit is generated by protein deiminase 4 (PAD4), which contributes to histone citrullination and chromatin decondensation during NETosis ^35^. We found no difference in Sytox Green signals from PMA stimulated or PAO1 infected neutrophils from C57BL/6 compared with *Gsdmd^-/-^* mice, although there was less NETosis in neutrophils infected with the Δ*pscD* injectosome or the Δ*fliC* flagellin mutant (**Figure 5C,D; Figure S4A, B)**.

In agreement with the Sytox data, we found that up to 80% neutrophils were H3Cit+ following stimulation with PMA or infection with *P. aeruginosa* strain PAO1; however, there was no significant difference in H3Cit binding between neutrophils from C57BL/6 and *Gsdmd^-/-^* mice (**Figure 5E,F, Figure S4C)**. There were also no differences in NET formation between neutrophils from C57BL/6 and *Gsdme^-/-^* mice (**Figure 5I, J)**.

Additional studies showed that PMA treated neutrophils incubated with LDC7559 produced significantly less ROS (**Figure S4D,E)**, which is consistent with its reported effect on NADPH oxidase ^32^. We also show a representative time course of NETosis in the presence of GSDMD inhibitors (**Figure S4F)**. *P. aeruginosa* Δ*pscD* induces ROS production by human neutrophils that is inhibited in strains expressing ExoS ADP ribosyl transferase ^17^. We repeated this finding using murine neutrophils and found that ROS production induced by PMA or Δ*pscD* was not dependent on GSDMD **(Figure S4G,H)**, and that in contrast to ROS-dependent NETosis induced by PMA, NETosis induced by PAO1 is ROS independent (**Figure S4I-K).** In contrast to our findings that PAO1 induced NETosis is ROS independent, Parker *et. al*. showed that DPI inhibited NET formation induced by PMA, Candida or PAO1 ^33^. They used an MOI of 10:1 PAO1 and quantified Sytox at 4h; however, we show no DPI inhibition at any time point after infection, whereas in the same experiments, DPI inhibited PMA induced NETosis (**Figure S4I-K**).

Overall, these observations clearly demonstrate that GSDMD and GSDME are not required for NETosis induced by either PMA or following infection with T3SS expressing *P. aeruginosa*.

## DISCUSSION

Bacterial T3SS effectors are efficiently injected into target eukaryotic cells where they can compromise the host immune response. For example, *Shigella flexneri* T3SS ubiquitin ligase IpaH7.8 selectively targets human GSDMB and GSDMD for proteasomal degradation, thereby inhibiting both GSDMB mediated bacterial killing and GSDMD mediated host cell lysis to protect the replicative environment of the bacteria ^34, 35^. *Pseudomonas, Salmonella*, and *Yersinia* T3SS and *Legionella* Type IV secretion activate IL-1β secretion and pyroptosis in macrophages primarily through recognition of needle structure proteins by NAIP1 and NAIP2 (humans have a single NAIP), for activation of the NLRC4 inflammasome ^36^. Also, *Yersinia YopJ* acetyltransferase promotes macrophage pro-IL-1β and GSDMD cleavage and pyroptosis ^37, 38^

In contrast to macrophages, there are relatively few reports on the role of neutrophils as a source of IL-1β during bacterial infection. In *Burkholderia* infected neutrophils, IL-1β secretion and pro-GSDMD cleavage by neutrophils requires caspase-1 or caspase-11, and selective deletion of caspase-11 in neutrophils results in impaired bacterial killing ^39^. Further, survival of mice infected with *YopJ* expressing *Yersinia* is dependent on GSDME rather than GSDMD, and GSDME is required for IL-1β secretion by neutrophils rather than macrophages ^19^. Consistent with that report, we found that *P. aeruginosa* induces pro-GSDME cleavage in neutrophils, though not in macrophages; however, IL-1β secretion was dependent on GSDMD and not GSDME. Further, while pro-GSDME is known to be cleaved by caspase-3 ^15^, we found that pro-caspase-3 was not cleaved in *P. aeruginosa* infected neutrophils. As we and others reported that serine proteases such as neutrophil elastase can cleave pro-GSDMD ^8, 21, 40^, it is possible that serine proteases also cleave pro-GSDME in neutrophils. In support of this concept, Granzyme directly cleaves GSDME to drive lytic tumor cell death, and TLR4-activated neutrophils upregulate expression of granzyme B as part of their extensive complement of serine proteases ^25, 41^.

The differences in host and bacterial mediators of IL-1β secretion between macrophages and neutrophils infected with *P. aeruginosa* are summarized in **Figure 6.** Pre-incubation with LPS induces expression of NLRP3 and pro-IL-1β in macrophages and neutrophils; however, following macrophage infection with T3SS expressing *P. aeruginosa*, the needle and translocon structures selectively activate NLRC4 (through NAIP1 and NAIP2), resulting in caspase-1 mediated IL-1β and GSDMD cleavage, plasma membrane pore formation, pyroptosis and secretion of bioactive IL-1β, which is consistent with earlier reports on the role of NLRC4 in *P. aeruginosa* infected macrophages ^26, 27, 42^. In contrast, we found that the needle structure can activate NLRC4 in neutrophils in the absence of exoenzymes (in Δ*exoST* mutants), leading to cleavage of GSDMD and IL-1β. However, in the presence of ExoS and ExoT, NLRP3 rather than NLRC4 was required for pro-GSDMD and IL-1β cleavage. We identified ExoS ADPRT as the mediator of selective NLRP3 activation. Importantly, we also demonstrate using *Nlrp3^-/-^*, *Nlrc4^-/-^* mice and the MCC950 NLRP3 inhibitor that NLRP3 rather than NLRC4 is required for bacterial killing and to prevent corneal perforation in *P. aeruginosa* keratitis.

**Figure 6.**
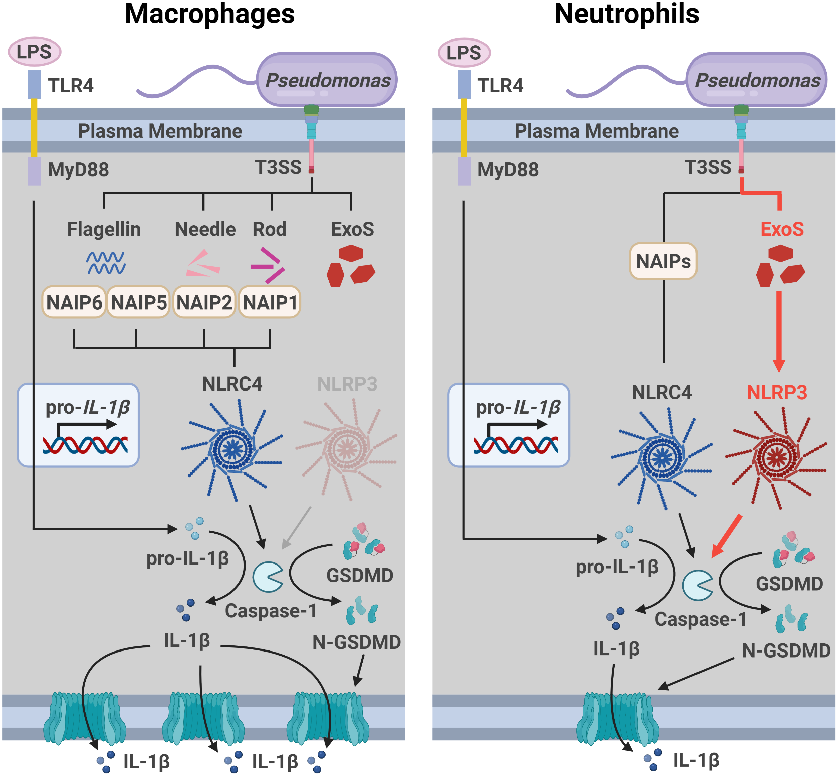
Predicted sequence of events in *P. aeruginosa* induced IL-1b secretion by macrophages and neutrophils. See text for explanation.

Collectively, these findings demonstrate a physiological role for NLRP3 and caspase-1 in regulating infection in a clinically relevant murine model of blinding corneal disease. In contrast to these observations, Santoni *et al*. reported that IL −1β secretion and pyroptosis in PAO1 infected neutrophils was strictly dependent on NLRC4 ^28^. Although we were able to reproduce their observations using bone marrow neutrophils that were not primed with LPS prior to infection, we reported that NLRP3 expression in neutrophils needs to be induced by TLR priming ^6^. In support of this, IL-1β secretion by casein - induced peritoneal neutrophils infected for 1h with PAO1 was completely dependent on NLRP3 in the absence of in vitro LPS priming due to the *in vivo* priming that occurs as recruited neutrophils accumulate in the inflammatory milieu of the peritoneum (no LPS priming). Also, in *P. aeruginosa* infection, NLRP3 and caspase-1, but not NLRC4 were required for IL-1β production, bacterial killing and corneal disease severity. While Santoni *et al* showed a role for caspase-1 in a *P. aeruginosa* lung infection model, they did not examine if there is a role for NLRC4 *in vivo ^28^*. We therefore conclude that selective usage of NLRP3 more closely resembles the activation state of neutrophils during infection and inflammation.

We also examined the role of GSDMD and GSDME during infection. Although GSDMD and GSDME were cleaved during corneal infection with PAO1, there was no effect of GSDMD and/or GSDME deficiency in on the progression of corneal disease or on bacterial killing. While we have yet to identify the underlying mechanisms, it is likely that it is due to GSDMD independent IL-1β secretion by neutrophils at later time points *in vitro* and in *Salmonella typhimurium* infected mice as described by Schroder and colleagues ^5^.

The underlying mechanisms that regulate inflammasome dependence between neutrophils and macrophages are unclear, although as macrophages express more ASC and caspase-1 and caspase-11 than neutrophils on a per cell basis ^43^, it is possible that macrophages also express higher levels of NAIPs or NLRC4 than neutrophils, resulting in more rapid NLRC4 mediated responses in macrophages. However, despite the *P. aeruginosa* needle structure activating NLRC4 in neutrophils, we found IL-1β secretion induced by ExoS expressing strains to be NLRP3 dependent. NLRP3 may also be a substrate for ExoS ADPRT, resulting in increased activity, similar to the gain-of-function mutations in NLRP3 that cause cryopyrin associated period syndromes (CAPS) ^44^. Neutrophils are the major source of IL-1β in CAPS patients, and gain-of-function mutations expressed only in neutrophils are sufficient to induce CAPS like symptoms in mice ^45^. NLRP3 activity is also regulated by post-translational modifications, including JNK phosphorylation of NLRP3, which potentiates oligomerization during LPS priming ^46^. Enzymes that regulate ubiquitylation and deubiquitylation also regulate NRLP3 activity ^47, 48^. Additionally, the *Mycoplasma pneumoniae* community-acquired respiratory distress syndrome (CARDS) toxin ADP ribosylates NLRP3 to enhance inflammasome activity ^49, 50^. In another study, macrophages incubated with recombinant CARDS toxin induced NLRP3-mediated IL-1β secretion via a mechanism requiring the ADPRT enzymatic activity of CARDS, and CARDS toxin interaction with NLRP3 ^50^, implying that ADP ribosylation resulted in increased NLRP3 activity. Similarly, Group A Streptococcus *SpyA* ribosyl transferase toxin induced NLRP3/caspase-1 dependent IL-1β secretion by macrophages, although these investigators did not examine inflammasome usage ^51,52^.

Finally, it will be relevant to consider how additional intracellular signals triggered by PAO1 infection of neutrophils might contribute to NLRP3 activation. Potential signals are increased K^+^ efflux via the translocon pores ^53, 54^ and accumulation of phosphatidylinositol-4-phosphate in the endosomal or dispersed trans-Golgi vesicles that recruit NLRP3 for subsequent trafficking to the NEK7-containing microtubule organizing centers where active NLRP3 inflammasomes are assembled ^19, 55, 56^.

An additional mechanism for the PAO1 induced redirection from NLRC4 to NLRP3 inflammasome usage is that ExoS ADPRT inhibits NAIP or NLRC4, which remain inactive even in the absence of NLRP3. Although most reports cite modifications that enhance NLRC4 activity ^53, 54^, ExoT was recently found to inhibit NLRC4 activation by blocking Crk2/PKCδ – mediated phosphorylation of NLRC4, which is required for activation ^57^. ExoS ADPRT modifications of NLRP3 and NLRC4 in neutrophils compared with macrophages will be examined in future studies. An important caveat in performing these experiments is that ExoS ADPRT is highly promiscuous compared with as it has multiple substrates compared with ExoT ADPRT ^10^. In support of this concept, a recent report showed that expression of ExoS ADPRT correlated with decreased lytic cell death in corneal epithelial cells ^58^, although the target proteins were not identified.

We reported that activation of the NLRP3 inflammasome in neutrophils by *Streptococcus pneumoniae* does not lead to pyroptotic cell death as quantified by LDH release ^6, 7^. Here we find that T3SS expressing *P. aeruginosa* induce release of low levels of LDH (~30% of triton control), compared with macrophages, suggesting that this level of neutrophil pyroptosis may not be physiologically significant.

In addition to its role in mediating secretion of IL-1β and pyroptotic death, GSDMD in neutrophils has been linked to the release of neutrophil extracellular traps (NETs) and NETosis, a mode of lytic cell death associated with many but not all stimuli that trigger NET release ^20, 21^. Conversely, more recent studies have questioned the requirement for GSDMD in NET release and NETosis ^28, 59^, which is in agreement with results in the current study wherein GSDMD or GSDME had no apparent effect on NETosis induced by PMA or *P. aeruginosa* when quantified by Sytox staining of released DNA or accumulation of H3Cit using *Gsdmd^-/-^* or *Gsdme^-/-^* neutrophils or C57/BL6 neutrophils treated with GSDMD small molecule inhibitors. This discrepancy may relate to recent findings by Santoni *et al* that although GSDMD is not required for *P. aeruginosa* induced NETosis, it appears to have a role in mediating DNA and histone release from the nucleus into the cytoplasm ^28^.

In an earlier report, we showed that PAO1 infection of peritoneal neutrophils induced release of IL-1β that was attenuated (~50-60%) but not completely suppressed in neutrophils from *caspase-1^-/-^* and *Nlrc4^-/-^* mice ^60^. However, production of mature IL-1β and bacterial killing in corneal infections were not inhibited in the absence of caspase-1 or NLRC4; rather, it was predominantly mediated by neutrophil elastase ^60^. In the current study, we repeated the infection studies in neutrophil elastase (*Elane*)^*-/-*^ mice and found no differences in CFU, corneal disease or in IL-1β cleavage. While this difference is puzzling, we suspect that it is related bacterial growth conditions that can have a profound effect on the ability of *P. aeruginosa* to produce and assemble the T3SS ^61^. Therefore, bacteria grown in high salt LB in the current study likely induced higher T3SS that is required to induce the NRLP3/caspase-1 pathway in neutrophils. In agreement, Santoni *et al* also found no role for neutrophil serine proteases in IL-1β secretion ^28^.

Overall, results from the current study reveal a previously unappreciated role for specific proteins of the bacterial T3SS in selectively directing activation of the NLRP3 versus NLRC4 inflammasomes in neutrophils but not macrophages. The findings also increase the understanding of how GSDMD versus GSDME cleavage is differentially utilized for IL-1β secretion and regulated cell death responses in neutrophils compared with macrophages, and in bacterial infections where neutrophils are the predominant cell type.

## Data availability

The authors declare that data supporting the findings of the current study are available within the article and Supplementary Information or available from the corresponding authors upon request.

## Author contributions

MSM, TL and KL designed and performed *in vitro* experiments and MSM, KL, SA, MEM, BM and JT performed the *in vivo* experiments. MM, TL, KL, GRD, AR, and EP wrote the manuscript.

## METHODS

**Reagents:** All reagents and antibodies are listed in Supplementary Table 1.

### Source of mice

C57BL/6, *gsdme^-/-^* and *nlrp3^-/-^* mice were purchased from Jackson Laboratories. *Gsdmd^-/-^* and *nlrc4^-/-^* mice were a kind gift from Dr. Russell Vance (University of California, Berkeley). *Gsdmd^-/-^ gsdme^-/-^* mice were generated in-house. All transgenic mice were on a C57BL/6 background. Animals were housed in pathogen free conditions in microisolator cages and were treated according to institutional guidelines following approval by the University of California IACUC.

### Bacteria

DMSO stocks of *P. aeruginosa* PAO1F and T3SS mutant strains were grown overnight in modified high salt LB (HSLB) broth containing 200 mM NaCl, 10 mM MgCl2, 0.5 mm CaCl_2_, and 5 mM EGTA to induce T3SS expression ^56^. For experiments, overnight cultures were diluted 1:100 in fresh HSLB broth and grown to early log phase, OD_600_ of 0.18-0.2. Bacteria were washed in PBS, resuspended at 3×10^8^ bacteria/mL, and 20uL containing 6x 10^6^ bacteria were added to 200,000 neutrophils (MOI of 30) for *in vitro* stimulations.

### Human blood neutrophils

Whole blood was collected from volunteers ages 18 and 65 years as part of the healthy blood donor program that is run by the Institute for Clinical and Translational Science at UC Irvine. This program has been approved by Institutional Review Board of the University of California (Irvine, CA), following informed consent, and donors are de-identified. Neutrophils were isolated using density gradient centrifugation (Ficoll-Paque Plus, GE Healthcare) at 500 × *g* for 30 min, which yields >90% purity as assessed by flow cytometry using anti-human CD16 and CD66b Abs (eBiosciences).

### Isolation of murine bone marrow and peritoneal neutrophils

Bone marrow neutrophils were collected from the tibias and femurs of immunocompetent C57BL/6 or gene knockout mice. Peritoneal neutrophils were elicited by injecting mice IP with 1 mL of a 9% sterile casein solution for 16-20h followed by a second injection of casein 3h prior to collection. Following lavage of the peritoneum, cells were drawn into cold PBS. Neutrophils were enriched via negative selection magnetic beads using the manufacturer’s recommended protocols (Biolegend MojoSort or STEMCELL EasySep),. This protocol routinely yields >95% neutrophils as determined by flow cytometry using antibodies to Ly6G and CD11b. Neutrophils were resuspended in RPMI (Gibco) at the specified concentrations for *in vitro* assays.

### Bone marrow derived macrophages (BMDM)

To derived macrophages, total bone marrow cells were collected by seeding total bone marrow cells into untreated culture flasks (Gen Clone, 25-214) containing 10 mL Dulbecco’s modified Eagle’s medium (DMEM; Gibco), 10% fetal bovine serum (FBS; Corning), and 10 ng/mL M-CSF (R&D Systems). On Day 2 and Day 4, non-adherent cells were removed by washing, and growth medium was replaced. On Day 6, BMDMs were detached using Cellstripper™ (Corning, NY) following manufacturer’s directions and plated in media with 20 ng/mL GM-CSF (STEMCELL) for *in vitro* assays the following day.

### Cell stimulation conditions

Isolated neutrophils and macrophages were stimulated with ultrapure lipopolysaccharide (LPS) (Invivogen) at 500 ng/mL for 3 h, and 3 mM ATP (Sigma-Aldrich) was added for 1 h to selectively activate NLRP3. To induce caspase-3 and GSDME activation, 100 ng/mL recombinant TNFα (R&D Systems) with 2 μM Birinapant Smac mimetic (MedChemExpress) or 2.5 ng/mL cycloheximide (final concentrations) were added for 4h. The pan caspase inhibitor Z-VAD-FMK (APExBIO), the NLRP3 inhibitor MCC950 (Invivogen) and the ROS inhibitor diphenyleneiodonium chloride (DPI, Sigma) were dissolved in dimethyl sulfoxide (DMSO) (Thermo Fisher Scientific). Cells were pretreated with ZVAD at 50 μM for 30 min, with MCC920 at 2 μM for 40 min, with DPI at 10 μM for 20 min, or with the equivalent volume of DMSO as vehicle control. Cells were incubated with *Pseudomonas aeruginosa* strains at an MOI=30 for 25 min for western blots or 1h for cytokine production and LDH release. Neutrophils were suspended in media containing 20 ng/mL GM-CSF to reduce spontaneous activation of Caspase-3. Macrophages were pretreated with 5 mM glycine for 30 min to inhibit macrophage pyroptosis as described ^25, 26^.

### Cytokine assays

Neutrophils (2×10^5^/well) or macrophages (3.5×10^4^ /well) were stimulated as indicated. Cell-free supernatants were collected by centrifugation in V-bottom 96-well plates (PlateOne). Proteins were measured by ELISA following manufacturer’s recommendations (DuoSet, R&D).

IL-1β bioactivity was quantified using the HEK-blue IL-1R1 reporter cell-line (InvivoGen). 2.8 ×10^5^ cells/mL in DMEM (GIBCO) complete media with or without anti-IL-1β neutralizing antibodies (R&D Systems) and incubated overnight at 37 C with 5%CO2. Supernatants were incubated with QUANTI-Blue (InvivoGen) for 30 minutes, and cytokines were measured on the Biotek Cytation-5 reader. IL-1 concentrations were calculated as pg/ml based on standards.

### Lactate dehydrogenase (LDH) assay for cell death

Cell cytotoxicity was determined by quantifying LDH release with CytoTox 96^®^ Non-Radioactive Cytotoxicity Assay (Promega) according to the manufacturer’s instructions. Percentage cytotoxicity was calculated based on maximum LDH release following incubation with 1% Triton X-100.

### Western Blot

Cells (4×10^6^ BMNs or 2×10^6^ BMDMs) were lysed in 1 x lysis buffer (CST). For neutrophils, we included 5 mM diisopropylfluorophosphate (DFP), which is a covalent inhibitor of neutral serine proteases such as neutrophil elastase. We found that DFP inhibits post-lysis processing of neutrophil proteins, including GSDMD ^7^. Protein in 1-1.5 mL of supernatants was precipitated using 10% sodium deoxycholate and 100% TCA. The resulting pellets were washed with 100% acetone and resuspended in 0.2 M NaOH. BCA assays were run to determine protein concentrations, and 10-30 μg total protein in 1-1.5 mL of TCA-precipitated supernatant was separated on 4-20% acrylamide gels via SDS-PAGE. Protein was transferred to PVDF membranes (Millipore) using the TransBlot Turbo semi-dry apparatus (Bio-Rad).

Antibodies for western blot were diluted as follows: anti-IL-1β(1:800), anti-β-actin (1:500), anti-GSDMD (1:500), anti-GSDME (1:1000), anti-caspase-3 (1:1000), HRP-conjugated secondary antibodies (1:1000). Membranes were developed using Supersignal West Femto Maximum Sensitivity Substrate (Thermo Scientific).

### Reactive oxygen species (ROS) quantification

Neutrophils were incubated with 50 μM luminol (Sigma) at 1×10^6^ cells/mL. Cells were plated in black-wall 96-well plates with an optically clear bottom (CoStar 3720) at 2×10^5^ neutrophils per well and incubated for 20-30 minutes. *P. aeruginosa* strains grown to log phase or other stimuli were added at indicated concentrations. Cells were incubated in the Cytation5 (BioTek) at 37°C and luminescence was measure from each well every 2 min for 90 min. Area under the curve was calculated using Prism (Graphpad).

### Neutrophil extracellular traps

#### Sytox^™^ kinetic assay for extracellular DNA

Enriched neutrophils were resuspended at 1×10^6^ cells/mL in RPMI with 20 ng/mL GM-CSF (R&D or STEMCELL) (for murine neutrophils) or RPMI with 2% FBS (for human neutrophils) with 1μM Sytox Green (Invitrogen) and plated in 96-well plates at 200,000 neutrophils per well. Log phase *P. aeruginosa* and other stimuli were added at indicated concentrations, and fluorescence (485 Ex/525 Em) was quantified every 15 min for 16h using a Cytation5 (BioTek) at 37°C with 5% CO2. The area under the curve between time 0-6h was calculated using Prism (Graphpad).

#### Histone citrullination

Enriched casein-elicited peritoneal mouse neutrophils were resuspended in RPMI with 20 ng/mL GM-CSF (R&D or STEMCELL) at 1×10^6^ cells/mL. 500,000 neutrophils were seeded onto sterile 12mm circular glass coverslips (#1.5, Thermo or EMS) in 24-well plates and stimulated as indicated. Coverslips were washed gently with cold PBS and fixed for 20 minutes at 4°C in 2% PFA in PBS. For washes and fixation, CaCl2 and MgCl2 were added to PBS at 0.9 mM and 0.5 mM, respectively. Coverslips were incubated for 1h in 10% normal donkey serum in PBS with 0.1% Triton X-100 and incubated overnight with indicated primary antibodies in blocking buffer. Coverslips were then washed in PBS, incubated in secondary antibodies for 2h, and counterstained in 0.1μg/mL DAPI before being mounted to slides using VectaSHIELD HardSet (Vector Labs) and sealed with nail polish. Neutrophils were imaged using an LSM700 confocal microscope (Zeiss) and processed with FIJI/ImageJ (NIH) or Zen 2 blue edition (Zeiss).

### *Murine model of P. aeruginosa* corneal infection

Overnight cultures of *P. aeruginosa* PAO1F or mutant strains were grown to log phase (OD_600_ of 0.2) in LB broth, then washed and resuspended in PBS at 5×10^7^ bacteria/ml. C57BL/6 and transgenic mice 7-12 weeks old were anesthetized with ketamine/xylazine solution, the corneal epithelium was abraded with three parallel scratches using a sterile 26-gauge needle, and 2 μL of a suspension of bacteria were added topically (approximately 1×10^5^ bacteria per eye).

After 24h or 48h, mice were euthanized and corneas were imaged by brightfield to detect opacification, or by fluorescence microscopy to detect GFP-expressing bacteria. Fluorescent intensity images were quantified using Image J software (NIH). For live bacteria, whole eyes were homogenized in PBS using a TissueLyser II (Qiagen, 30 Hz for 3 minutes), and homogenates were serially diluted and streaked on LB plates for quantification of colony forming units (CFU) by manual counting. CFU were also determined at 2 h to confirm the inoculum. For western blot and cytokine analyses, corneas were carefully dissected to remove vascularized iris tissue, and placed in 1x lysis buffer (CST) containing 5 mM DFP, and processed as described above.

### Flow cytometry

Dissected corneas were incubated with 3 mg/ml collagenase (C0130; Sigma-Aldrich) in RPMI (Life Technologies), with 1% HEPES (Life Technologies), 1% penicillin-streptomycin (Life Technologies), and 0.5% BSA (Fisher Bioreagents) for 1 h and 15 min at 37·C. Cells were incubated for 5 min with anti-mouse CD16/32 Ab (BioLegend) to block Fc receptors. Then cells were incubated 20 min at 4·C with anti-mouse CD45-allophycocyanin, Ly6GBV510, Ly6C-PE-Cy7, CD11b-PETxRed, CCR2-BV421, and F4/80-FITC (BioLegend) and fixable viability dye (BD Biosciences). Cells were washed with FACS buffer and quantified using an ACEA Novocyte flow cytometer and NovoExpress software. Neutrophils were identified as CD45+, CD11b+, Ly6G+, CCR2-, and monocytes were CD45+, CD11b+, Ly6G-CCR2+.

### Statistics

Statistical significance was determined using unpaired t-test or two-way ANOVA with either HSD Tukey’s post hoc analysis (GraphPad Prism). Differences were considered significant when the *P* value was <0.05.

## References

1. Liu X, Xia S, Zhang Z, Wu H, Lieberman J. Channelling inflammation: gasdermins in physiology and disease. Nat Rev Drug Discov 20, 384–405 (2021).

2. Karmakar M, Sun Y, Hise AG, Rietsch A, Pearlman E. Cutting edge: IL-1beta processing during Pseudomonas aeruginosa infection is mediated by neutrophil serine proteases and is independent of NLRC4 and caspase-1. J Immunol 189, 4231–4235 (2012).

3. Karmakar M, Katsnelson MA, Dubyak GR, Pearlman E. Neutrophil P2X7 receptors mediate NLRP3 inflammasome-dependent IL-1beta secretion in response to ATP. Nat Commun 7, 10555 (2016).

4. Karmakar M, et al. Neutrophil IL-1beta processing induced by pneumolysin is mediated by the NLRP3/ASC inflammasome and caspase-1 activation and is dependent on K+ efflux. J Immunol 194, 1763–1775 (2015).

5. Chen KW, et al. The neutrophil NLRC4 inflammasome selectively promotes IL-1beta maturation without pyroptosis during acute Salmonella challenge. Cell Rep 8, 570–582 (2014).

6. Monteleone M, et al. Interleukin-1beta Maturation Triggers Its Relocation to the Plasma Membrane for Gasdermin-D-Dependent and -Independent Secretion. Cell Rep 24, 1425–1433 (2018).

7. Karmakar M, et al. N-GSDMD trafficking to neutrophil organelles facilitates IL-1beta release independently of plasma membrane pores and pyroptosis. Nat Commun 11, 2212 (2020).

8. Hauser AR, The type III secretion system of Pseudomonas aeruginosa: infection by injection. Nat Rev Microbiol 7, 654–665 (2009).

9. Munder A, et al. The Pseudomonas aeruginosa ExoY phenotype of high-copy-number recombinants is not detectable in natural isolates. Open Biol 8, (2018).

10. Ung L, Bispo PJM, Shanbhag SS, Gilmore MS, Chodosh J. The persistent dilemma of microbial keratitis: Global burden, diagnosis, and antimicrobial resistance. Surv Ophthalmol 64, 255–271 (2019).

11. Karthikeyan RS, et al. Host response and bacterial virulence factor expression in Pseudomonas aeruginosa and Streptococcus pneumoniae corneal ulcers. PLoS One 8, e64867 (2013).

12. Borkar DS, et al. Association between cytotoxic and invasive Pseudomonas aeruginosa and clinical outcomes in bacterial keratitis. JAMA Ophthalmol 131, 147–153 (2013).

13. Sun Y, Karmakar M, Taylor PR, Rietsch A, Pearlman E. ExoS and ExoT ADP ribosyltransferase activities mediate Pseudomonas aeruginosa keratitis by promoting neutrophil apoptosis and bacterial survival. J Immunol 188, 1884–1895 (2012).

14. Vareechon C, Zmina SE, Karmakar M, Pearlman E, Rietsch A. Pseudomonas aeruginosa Effector ExoS Inhibits ROS Production in Human Neutrophils. Cell Host Microbe 21, 611–618e615 (2017).

15. Broz P, Pelegrin P, Shao F. The gasdermins, a protein family executing cell death and inflammation. Nat Rev Immunol 20, 143–157 (2020).

16. Kambara H, et al. Gasdermin D Exerts Anti-inflammatory Effects by Promoting Neutrophil Death. Cell Rep 22, 2924–2936 (2018).

17. Sollberger G, et al. Gasdermin D plays a vital role in the generation of neutrophil extracellular traps. Sci Immunol 3, (2018).

18. Chen KW, et al. RIPK1 activates distinct gasdermins in macrophages and neutrophils upon pathogen blockade of innate immune signaling. Proc Natl Acad Sci U S A 118, (2021).

19. Miao EA, et al. Innate immune detection of the type III secretion apparatus through the NLRC4 inflammasome. Proc Natl Acad Sci U S A 107, 3076–3080 (2010).

20. Chen KW, et al. Noncanonical inflammasome signaling elicits gasdermin D-dependent neutrophil extracellular traps. Sci Immunol 3, (2018).

21. Zhao Y, et al. The NLRC4 inflammasome receptors for bacterial flagellin and type III secretion apparatus. Nature 477, 596–600 (2011).

22. Zhao Y, Shao F. The NAIP-NLRC4 inflammasome in innate immune detection of bacterial flagellin and type III secretion apparatus. Immunol Rev 265, 85–102 (2015).

23. Vance RE. The NAIP/NLRC4 inflammasomes. Curr Opin Immunol 32, 84–89 (2015).

24. Sutterwala FS, Mijares LA, Li L, Ogura Y, Kazmierczak BI, Flavell RA. Immune recognition of Pseudomonas aeruginosa mediated by the IPAF/NLRC4 inflammasome. J Exp Med 204, 3235–3245 (2007).

25. Evavold CL, Ruan J, Tan Y, Xia S, Wu H, Kagan JC. The Pore-Forming Protein Gasdermin D Regulates Interleukin-1 Secretion from Living Macrophages. Immunity 48, 35–44e36 (2018).

26. Loomis WP, den Hartigh AB, Cookson BT, Fink SL. Diverse small molecules prevent macrophage lysis during pyroptosis. Cell Death Dis 10, 326 (2019).

27. Coll RC, et al. A small-molecule inhibitor of the NLRP3 inflammasome for the treatment of inflammatory diseases. Nat Med 21, 248–255 (2015).

28. Fuchs TA, et al. Novel cell death program leads to neutrophil extracellular traps. J Cell Biol 176, 231–241 (2007).

29. Brinkmann V, et al. Neutrophil extracellular traps kill bacteria. Science 303, 1532–1535 (2004).

30. Skopelja-Gardner S, et al. Regulation of Pseudomonas aeruginosa-Mediated Neutrophil Extracellular Traps. Front Immunol 10, 1670 (2019).

31. Thanabalasuriar A, et al. Neutrophil Extracellular Traps Confine Pseudomonas aeruginosa Ocular Biofilms and Restrict Brain Invasion. Cell Host Microbe 25, 526–536e524 (2019).

32. Rathkey JK, et al. Chemical disruption of the pyroptotic pore-forming protein gasdermin D inhibits inflammatory cell death and sepsis. Sci Immunol 3, (2018).

33. Hu JJ, et al. FDA-approved disulfiram inhibits pyroptosis by blocking gasdermin D pore formation. Nat Immunol 21, 736–745 (2020).

34. Amara N, et al. Selective activation of PFKL suppresses the phagocytic oxidative burst. Cell 184, 4480–4494e4415 (2021).

35. Sollberger G, Tilley DO, Zychlinsky A. Neutrophil Extracellular Traps: The Biology of Chromatin Externalization. Dev Cell 44, 542–553 (2018).

36. Sun Y, et al. TLR4 and TLR5 on corneal macrophages regulate Pseudomonas aeruginosa keratitis by signaling through MyD88-dependent and -independent pathways. J Immunol 185, 4272–4283 (2010).

37. Hansen JM, et al. Pathogenic ubiquitination of GSDMB inhibits NK cell bactericidal functions. Cell 184, 3178–3191e3118 (2021).

38. Luchetti G, et al. Shigella ubiquitin ligase IpaH7.8 targets gasdermin D for degradation to prevent pyroptosis and enable infection. Cell Host Microbe 29, 1521–1530e1510 (2021).

39. Bauer R, Rauch I. The NAIP/NLRC4 inflammasome in infection and pathology. Mol Aspects Med 76, 100863 (2020).

40. Sarhan J, et al. Caspase-8 induces cleavage of gasdermin D to elicit pyroptosis during Yersinia infection. Proc Natl Acad Sci U S A 115, E10888–E10897 (2018).

41. Orning P, et al. Pathogen blockade of TAK1 triggers caspase-8-dependent cleavage of gasdermin D and cell death. Science 362, 1064–1069 (2018).

42. Kovacs SB, et al. Neutrophil Caspase-11 Is Essential to Defend against a Cytosol-Invasive Bacterium. Cell Rep 32, 107967 (2020).

43. Franchi L, Stoolman J, Kanneganti TD, Verma A, Ramphal R, Nunez G. Critical role for Ipaf in Pseudomonas aeruginosa-induced caspase-1 activation. Eur J Immunol 37, 3030–3039 (2007).

44. Boucher D, et al. Caspase-1 self-cleavage is an intrinsic mechanism to terminate inflammasome activity. J Exp Med 215, 827–840 (2018).

45. Booshehri LM, Hoffman HM. CAPS and NLRP3. J Clin Immunol 39, 277–286 (2019).

46. Stackowicz J, et al. Neutrophil-specific gain-of-function mutations in Nlrp3 promote development of cryopyrin-associated periodic syndrome. J Exp Med 218, (2021).

47. Song N, et al. NLRP3 Phosphorylation Is an Essential Priming Event for Inflammasome Activation. Mol Cell 68, 185–197e186 (2017).

48. Liang Z, Damianou A, Di Daniel E, Kessler BM. Inflammasome activation controlled by the interplay between post-translational modifications: emerging drug target opportunities. Cell Commun Signal 19, 23 (2021).

49. Zangiabadi S, Abdul-Sater AA. Regulation of the NLRP3 Inflammasome by Posttranslational Modifications. J Immunol 208, 286–292 (2022).

50. Bose S, Segovia JA, Somarajan SR, Chang TH, Kannan TR, Baseman JB. ADP-ribosylation of NLRP3 by Mycoplasma pneumoniae CARDS toxin regulates inflammasome activity. mBio 5, (2014).

51. Segovia JA, et al. NLRP3 Is a Critical Regulator of Inflammation and Innate Immune Cell Response during Mycoplasma pneumoniae Infection. Infect Immun 86, (2018).

52. Lin AE, et al. A group A Streptococcus ADP-ribosyltransferase toxin stimulates a protective interleukin 1beta-dependent macrophage immune response. mBio 6, e00133 (2015).

53. Bardet J, et al. NLRC4 GOF Mutations, a Challenging Diagnosis from Neonatal Age to Adulthood. J Clin Med 10, (2021).

54. Canna SW, et al. An activating NLRC4 inflammasome mutation causes autoinflammation with recurrent macrophage activation syndrome. Nat Genet 46, 1140–1146 (2014).

55. Kroken AR, et al. Exotoxin S secreted by internalized Pseudomonas aeruginosa delays lytic host cell death. PLoS Pathog 18, e1010306 (2022).

56. Rietsch A, Mekalanos JJ. Metabolic regulation of type III secretion gene expression in Pseudomonas aeruginosa. Mol Microbiol 59, 807–820 (2006).

## References

1. Broz P, Pelegrin P, Shao F. The gasdermins, a protein family executing cell death and inflammation. Nat Rev Immunol 20, 143–157 (2020).

2. Liu X, Xia S, Zhang Z, Wu H, Lieberman J. Channelling inflammation: gasdermins in physiology and disease. Nat Rev Drug Discov 20, 384–405 (2021).

3. Fu J, Wu H. Structural Mechanisms of NLRP3 Inflammasome Assembly and Activation. Annu Rev Immunol, (2022).

4. Chen KW, et al. The neutrophil NLRC4 inflammasome selectively promotes IL-1beta maturation without pyroptosis during acute Salmonella challenge. Cell Rep 8, 570–582 (2014).

5. Monteleone M, et al. Interleukin-1beta Maturation Triggers Its Relocation to the Plasma Membrane for Gasdermin-D-Dependent and -Independent Secretion. Cell Rep 24, 1425–1433 (2018).

6. Karmakar M, et al. Neutrophil IL-1beta processing induced by pneumolysin is mediated by the NLRP3/ASC inflammasome and caspase-1 activation and is dependent on K+ efflux. J Immunol 194, 1763–1775 (2015).

7. Karmakar M, Katsnelson MA, Dubyak GR, Pearlman E. Neutrophil P2X7 receptors mediate NLRP3 inflammasome-dependent IL-1beta secretion in response to ATP. Nat Commun 7, 10555-(2016).

8. Karmakar M, et al. N-GSDMD trafficking to neutrophil organelles facilitates IL-1beta release independently of plasma membrane pores and pyroptosis. Nat Commun 11, 2212 (2020).

9. Dubyak GR, Miller BA, Pearlman E. Pyroptosis in neutrophils: Multimodal integration of inflammasome and regulated cell death signaling pathways. Immunol Rev, (2023).

10. Hauser AR. The type III secretion system of Pseudomonas aeruginosa: infection by injection. Nat Rev Microbiol 7, 654–665 (2009).

11. Jouault A, Saliba AM, Touqui L. Modulation of the immune response by the Pseudomonas aeruginosa type-III secretion system. Front Cell Infect Microbiol 12, 1064010 (2022).

12. Ung L, Bispo PJM, Shanbhag SS, Gilmore MS, Chodosh J. The persistent dilemma of microbial keratitis: Global burden, diagnosis, and antimicrobial resistance. Surv Ophthalmol 64, 255–271 (2019).

13. Cabrera-Aguas M, Khoo P, Watson SL. Infectious keratitis: A review. Clin Exp Ophthalmol 50, 543–562 (2022).

14. Borkar DS, et al. Association between cytotoxic and invasive Pseudomonas aeruginosa and clinical outcomes in bacterial keratitis. JAMA Ophthalmol 131, 147–153 (2013).

15. Karthikeyan RS, et al. Host response and bacterial virulence factor expression in Pseudomonas aeruginosa and Streptococcus pneumoniae corneal ulcers. PLoS One 8, e64867 (2013).

16. Sun Y, Karmakar M, Taylor PR, Rietsch A, Pearlman E. ExoS and ExoT ADP ribosyltransferase activities mediate Pseudomonas aeruginosa keratitis by promoting neutrophil apoptosis and bacterial survival. J Immunol 188, 1884–1895 (2012).

17. Vareechon C, Zmina SE, Karmakar M, Pearlman E, Rietsch A. Pseudomonas aeruginosa Effector ExoS Inhibits ROS Production in Human Neutrophils. Cell Host Microbe 21, 611–618e615 (2017).

18. Dubyak GR, Miller BA, Pearlman E. Pyroptosis in Neutrophils: Multimodal Integration of Inflammasome and Regulated Cell Death Signaling Pathways. Immunological Reviews In press, (2023).

19. Chen KW, et al. RIPK1 activates distinct gasdermins in macrophages and neutrophils upon pathogen blockade of innate immune signaling. Proc Natl Acad Sci U S A 118, (2021).

21. Sollberger G, et al. Gasdermin D plays a vital role in the generation of neutrophil extracellular traps. Sci Immunol 3, (2018).

23. Zhao Y, et al. The NLRC4 inflammasome receptors for bacterial flagellin and type III secretion apparatus. Nature 477, 596–600 (2011).

24. Vance RE. The NAIP/NLRC4 inflammasomes. Curr Opin Immunol 32, 84–89 (2015).

25. Zhang Z, et al. Gasdermin E suppresses tumour growth by activating anti-tumour immunity. Nature 579, 415–420 (2020).

26. Miao EA, et al. Innate immune detection of the type III secretion apparatus through the NLRC4 inflammasome. Proc Natl Acad Sci US A 107, 3076–3080 (2010).

27. Sutterwala FS, Mijares LA, Li L, Ogura Y, Kazmierczak BI, Flavell RA. Immune recognition of Pseudomonas aeruginosa mediated by the IPAF/NLRC4 inflammasome. J Exp Med 204, 3235–3245 (2007).

28. Santoni K, et al. Caspase-1-driven neutrophil pyroptosis and its role in host susceptibility to Pseudomonas aeruginosa. PLoS Pathog 18, e1010305 (2022).

29. Ratitong B, Marshall ME, Dragan MA, Anunciado CM, Abbondante S, Pearlman E. Differential Roles for IL-1alpha and IL-1beta in Pseudomonas aeruginosa Corneal Infection. J Immunol, (2022).

30. Brinkmann V, et al. Neutrophil extracellular traps kill bacteria. Science 303, 1532–1535 (2004).

31. Fuchs TA, et al. Novel cell death program leads to neutrophil extracellular traps. J Cell Biol 176, 231–241 (2007).

32. Amara N, et al. Selective activation of PFKL suppresses the phagocytic oxidative burst. Cell 184, 4480–4494e4415 (2021).

33. Parker HA, Jones HM, Kaldor CD, Hampton MB, Winterbourn CC. Neutrophil NET Formation with Microbial Stimuli Requires Late Stage NADPH Oxidase Activity. Antioxidants (Basel) 10, (2021).

34. Hansen JM, et al. Pathogenic ubiquitination of GSDMB inhibits NK cell bactericidal functions. Cell 184, 3178–3191e3118 (2021).

35. Luchetti G, et al. Shigella ubiquitin ligase IpaH7.8 targets gasdermin D for degradation to prevent pyroptosis and enable infection. Cell Host Microbe 29, 1521–1530e1510 (2021).

36. Bauer R, Rauch I. The NAIP/NLRC4 inflammasome in infection and pathology. Mol Aspects Med 76, 100863 (2020).

37. Sarhan J, et al. Caspase-8 induces cleavage of gasdermin D to elicit pyroptosis during Yersinia infection. Proc Natl Acad Sci U S A 115, E10888–E10897 (2018).

38. Orning P, et al. Pathogen blockade of TAK1 triggers caspase-8-dependent cleavage of gasdermin D and cell death. Science 362, 1064–1069 (2018).

39. Kovacs SB, et al. Neutrophil Caspase-11 Is Essential to Defend against a Cytosol-Invasive Bacterium. Cell Rep 32, 107967 (2020).

40. Kambara H, et al. Gasdermin D Exerts Anti-inflammatory Effects by Promoting Neutrophil Death. Cell Rep 22, 2924–2936 (2018).

41. Martin A, et al. Tumor-derived granzyme B-expressing neutrophils acquire antitumor potential after lipid A treatment. Oncotarget 9, 28364–28378 (2018).

42. Franchi L, Stoolman J, Kanneganti TD, Verma A, Ramphal R, Nunez G. Critical role for Ipaf in Pseudomonas aeruginosa-induced caspase-1 activation. Eur J Immunol 37, 3030-3039 (2007).

43. Boucher D, et al. Caspase-1 self-cleavage is an intrinsic mechanism to terminate inflammasome activity. J Exp Med 215, 827–840 (2018).

44. Booshehri LM, Hoffman HM. CAPS and NLRP3. J Clin Immunol 39, 277–286 (2019).

45. Stackowicz J, et al. Neutrophil-specific gain-of-function mutations in Nlrp3 promote development of cryopyrin-associated periodic syndrome. J Exp Med 218, (2021).

46. Song N, et al. NLRP3 Phosphorylation Is an Essential Priming Event for Inflammasome Activation. Mol Cell 68, 185–197e186 (2017).

47. Liang Z, Damianou A, Di Daniel E, Kessler BM. Inflammasome activation controlled by the interplay between post-translational modifications: emerging drug target opportunities. Cell Commun Signal 19, 23 (2021).

48. Zangiabadi S, Abdul-Sater AA. Regulation of the NLRP3 Inflammasome by Posttranslational Modifications. J Immunol 208, 286–292 (2022).

49. Bose S, Segovia JA, Somarajan SR, Chang TH, Kannan TR, Baseman JB. ADP-ribosylation of NLRP3 by Mycoplasma pneumoniae CARDS toxin regulates inflammasome activity. mBio 5, (2014).

50. Segovia JA, et al. NLRP3 Is a Critical Regulator of Inflammation and Innate Immune Cell Response during Mycoplasma pneumoniae Infection. Infect Immun 86, (2018).

51. Lin AE, et al. A group A Streptococcus ADP-ribosyltransferase toxin stimulates a protective interleukin 1beta-dependent macrophage immune response. mBio 6, e00133 (2015).

52. Richter J, et al. Streptolysins are the primary inflammasome activators in macrophages during Streptococcus pyogenes infection. Immunol Cell Biol 99, 1040–1052 (2021).

53. Kundracik E, Trichka J, Diaz Aponte J, Roistacher A, Rietsch A. PopB-PcrV Interactions Are Essential for Pore Formation in the Pseudomonas aeruginosa Type III Secretion System Translocon. mBio 13, e0238122 (2022).

54. Dortet L, Lombardi C, Cretin F, Dessen A, Filloux A. Pore-forming activity of the Pseudomonas aeruginosa type III secretion system translocon alters the host epigenome. Nat Microbiol 3, 378–386 (2018).

55. Zhang Z, et al. Distinct changes in endosomal composition promote NLRP3 inflammasome activation. Nat Immunol 24, 30–41 (2023).

56. Springer TI, Reid TE, Gies SL, Feix JB. Interactions of the effector ExoU from Pseudomonas aeruginosa with short-chain phosphatidylinositides provide insights into ExoU targeting to host membranes. J Biol Chem 294, 19012–19021 (2019).

57. Mohamed MF, et al. CrkII/Abl phosphorylation cascade is critical for NLRC4 inflammasome activity and is blocked by Pseudomonas aeruginosa ExoT. Nat Commun 13, 1295 (2022).

58. Kroken AR, et al. Exotoxin S secreted by internalized Pseudomonas aeruginosa delays lytic host cell death. PLoS Pathog 18, e1010306 (2022).

59. Chauhan D, et al. GSDMD drives canonical inflammasome-induced neutrophil pyroptosis and is dispensable for NETosis. EMBO Rep 23, e54277 (2022).

60. Karmakar M, Sun Y, Hise AG, Rietsch A, Pearlman E. Cutting edge: IL-1beta processing during Pseudomonas aeruginosa infection is mediated by neutrophil serine proteases and is independent of NLRC4 and caspase-1. J Immunol 189, 4231–4235 (2012).

61. Rietsch A, Mekalanos JJ. Metabolic regulation of type III secretion gene expression in Pseudomonas aeruginosa. Mol Microbiol 59, 807–820 (2006).

