## Supplemental Figures for "The NLRP3 inflammasome selectively drives IL-1β secretion by *Pseudomonas aeruginosa* infected neutrophils and regulates bacterial killing *in vivo*"

### LEGENDS TO SUPPLEMENTAL FIGURES

**Figure S1. IL-1β secretion and pyroptosis in peritoneal neutrophils.** Neutrophils were isolated from the peritoneal cavity and purified by negative bead selection following intraperitoneal injection of casein to induce sterile inflammation. **A,C.** IL-1β secretion; **B,D.** LDH release (**B**) following 1h infection with PAO1 or mutants in T3SS or flagellin (Δ*fliC). Peritoneal neutrophils were not primed with LPS.* Data points represent biological replicates from 2-3 repeat experiments.
**E. Caspase-8 and IL-1β cleavage in bone marrow neutrophils and bone marrow derived macrophages.**

**Figure S2. MCC950 inhibition of IL-1β secretion by C57BL/6 and *Nlrc4$^{-/-}$* bone marrow neutrophils. A, B.** Neutrophils isolated from C57BL/6 and *Nlrc4$^{-/-}$* mice were LPS primed for 3h and infected at a multiplicity of infection (MOI) of 30:1 bacteria: neutrophils with PAO1 or mutants for 1h in the presence of MCC950 or DMSO. IL-1β and LDH in culture supernatants were quantified. **C,D.** IL-1β and LDH in C57BL/6 and *Nlrc4$^{-/-}$* mice neutrophils that were either LPS primed and infected with PAO1 at an MOI of 10:1, or not primed and infected with PAO1 for 3h as described by Santoni *et. al.*, *PLoS Pathogens 2022*[1]. **E, F.** IL-1β and LDH in peritoneal neutrophils from C57BL/6 mice that were not LPS primed but were infected with 30:1 or 10:1 MOI for 1h in the presence of MCC950. Data points represent biological replicates from 2-3 repeat experiments.

**Figure S3.**
**A. Bioactive IL-1β in infected corneas.** IL-1R1 HEK293 cells from InVivogen incubated with homogenates from infected corneas in the presence of neutralizing antibodies to IL-1β.
**B-D. Neutrophil infiltration to PAO1 infected corneas. B-DC.** Neutrophils in C57BL/6 and *Nlrp3$^{-/-}$* corneas 24h post infection with PAO1 following collagenase digestion**. B,C.** representative flow cytometry scatter plots; **C.** quantification of total neutrophils. **E-G.** Neutrophils in C57BL/6 and *Gsdmd$^{-/-}$* corneas 24h post infection. **H.** IL-1β in homogenates from infected corneas measured by ELISA.
**I-K. Corneal infection in neutrophil elastase (*Elane*) gene knockout mice. G.** representative images of corneas of C57BL/6, *caspase-1,11$^{-/-}$,* and *Elane$^{-/-}$* mice were infected with PAO1 expressing green fluorescent protein (GFP). GFP expression **H** and CFU (**I**) showing significant difference between C57BL/6 and *caspase-1,11$^{-/-}$,* but not between C57BL/6 and *Elane$^{-/-}$* mice. Data points are biological replicates.

**Figure S4.**
**A-F. The role of GSDMD in neutrophil extracellular trap formation induced by PMA or infection with PAO1.**
**A-C. Effect of reported GSDMD inhibitors. A.** Representative time course of NETosis in PAO1 infected peritoneal neutrophils incubated with GSDMD inhibitors necrosulfonamide (NSA), disulfiram or LDC7559. **B,C.** Inhibition of ROS production by PMA – stimulated neutrophils incubated with DPI or by LDC7559 as described by Dixit in *Cell, 2021*[2]*.* B. representative time course; **B.** quantification of area under the curve. **C.** Data points are biological replicates.
**D, E.** Representative time course and combines data of NETosis induced by PAO1 compared with Δ*pscD* and Δ*fliC* mutants*.*
**F.** Representative images of H3Cit in PMA and PAO1 infected peritoneal neutrophils

**G-I.** Role of ROS in PMA stimulated, but not PAO1 infected peritoneal neutrophils. **G, H:** representative time courses; I: quantitative data

**J,K.** ROS production measured by Luminol in bone marrow neutrophils from C57BL/6 and *gsdmd-/-* mice following stimulation with PMA or infection with Δ*pscD* or PAO1. Representative time course (**J**) and combined data from biological replicates (**K**). Δ*pscD* induced higher ROS than PAO1 as we reported (*Cell Host & Microbe, 2017*[3]), and there was no difference between C57BL/6 and *gsdmd-/-* neutrophils.


1. Santoni K*, et al.* Caspase-1-driven neutrophil pyroptosis and its role in host susceptibility to Pseudomonas aeruginosa. *PLoS Pathog* **18**, e1010305 (2022).

2. Amara N*, et al.* Selective activation of PFKL suppresses the phagocytic oxidative burst. *Cell* **184**, 4480-4494 e4415 (2021).

3. Vareechon C, Zmina SE, Karmakar M, Pearlman E, Rietsch A. Pseudomonas aeruginosa Effector ExoS Inhibits ROS Production in Human Neutrophils. *Cell Host Microbe* **21**, 611-618 e615 (2017).

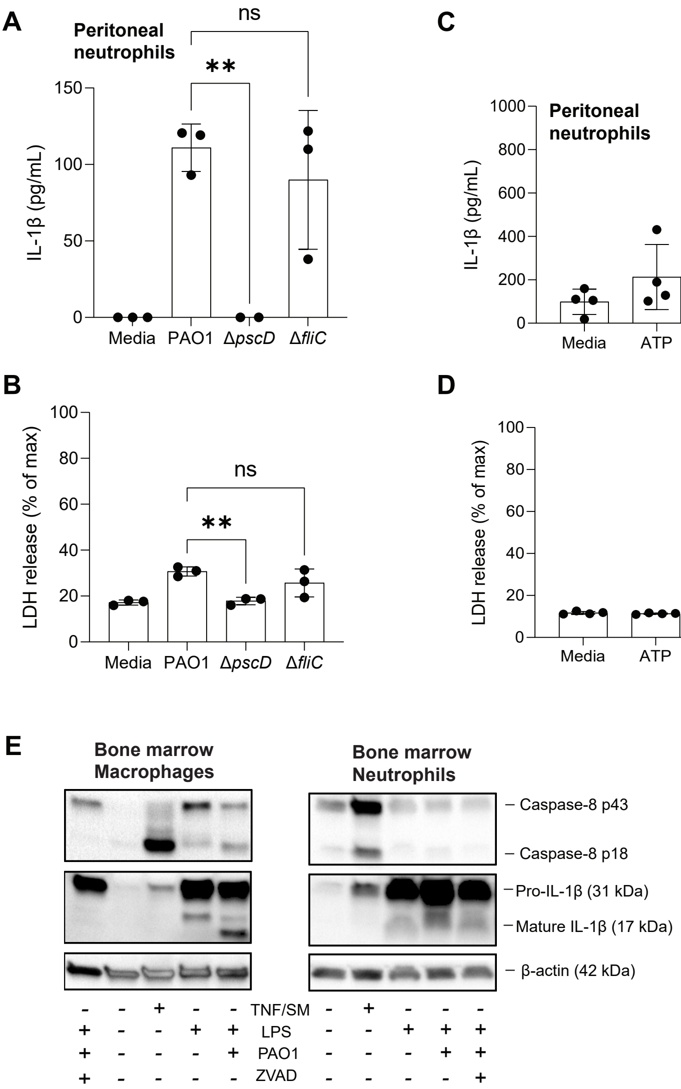
A
Peritoneal neutrophils
IL-1β (pg/mL)
Media
PAO1
ΔpscD
ΔfliC
ns
**
B
LDH release (% of max)
C
ATP
ΔpopB
ΔexoSTY
ΔexoST
D
****
E
Bone marrow Macrophages
Bone marrow Neutrophils
Caspase-8 p43
Caspase-8 p18
Pro-IL-1β (31 kDa)
Mature IL-1β (17 kDa)
β-actin (42 kDa)
TNF/SM
LPS
PAO1
ZVAD

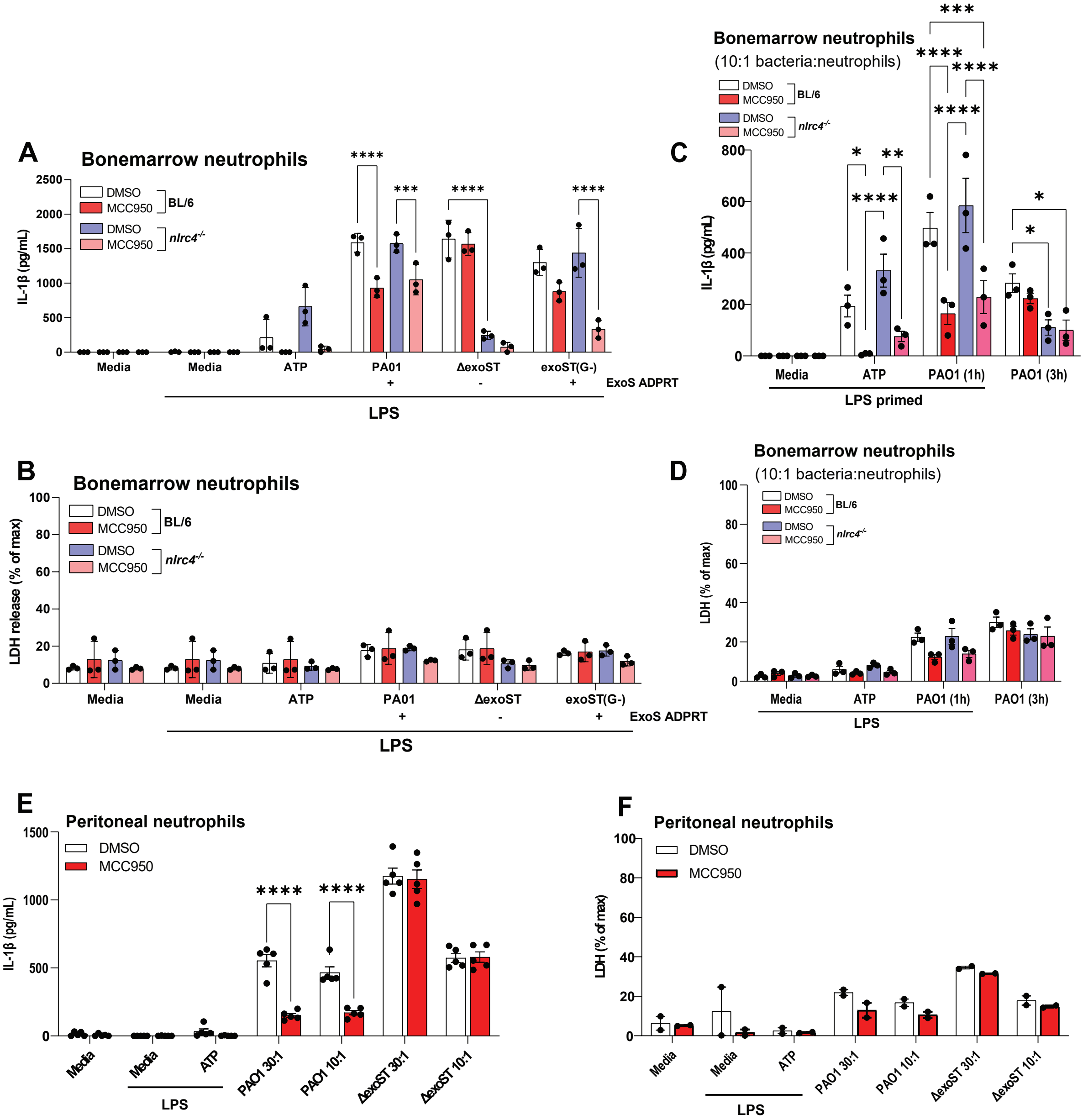

A
Bonemarrow neutrophils
IL-1β (pg/mL)
DMSO
MCC950
BL/6
DMSO
MCC950
nlrc4-/-
Media
Media
ATP
PA01
ΔexoST
exoST(G-)
+
-
+
ExoS ADPRT
LPS
B
Bonemarrow neutrophils
LDH release (% of max)
C
Bonemarrow neutrophils
(10:1 bacteria:neutrophils)
IL-1β (pg/mL)
Media
ATP
PAO1 (1h)
PAO1 (3h)
LPS primed
D
Bonemarrow neutrophils
(10:1 bacteria:neutrophils)
LDH (% of max)
LPS
E
Peritoneal neutrophils
IL-1β (pg/mL)
DMSO
MCC950
PAO1 30:1
PAO1 10:1
ΔexoST 30:1
ΔexoST 10:1
F
Peritoneal neutrophils
LDH (% of max)

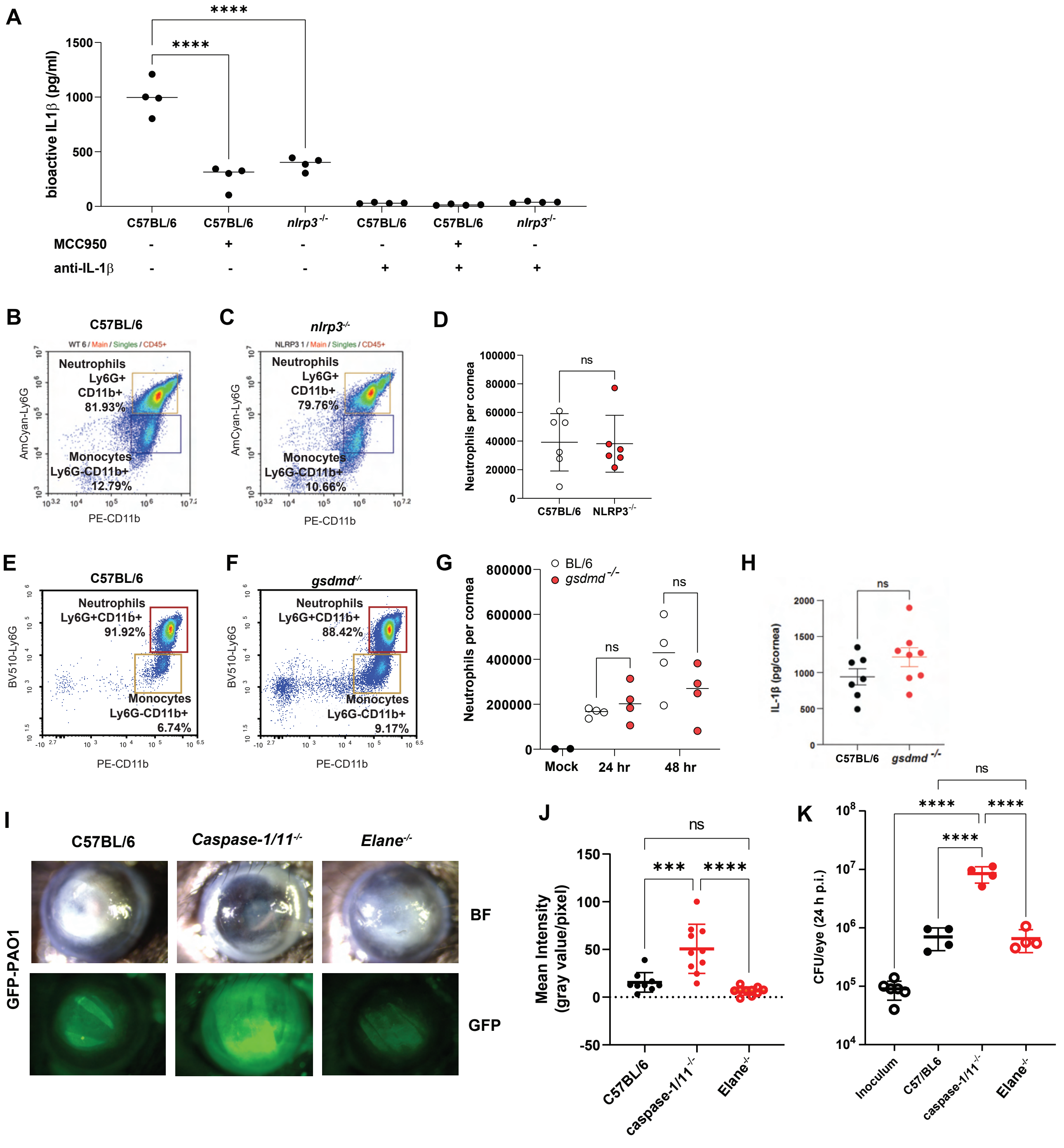
A
bioactive IL1β (pg/ml)
****
****
C57BL/6
C57BL/6
nlrp3-/-
C57BL/6
C57BL/6
nlrp3-/-
MCC950
anti-IL-1β
B
C57BL/6
Neutrophils Ly6G+ CD11b+ 81.93%
Monocytes Ly6G-CD11b+ 12.79%
AmCyan-Ly6G
PE-CD11b
C
nlrp3-/-
Neutrophils Ly6G+ CD11b+ 79.76%
Monocytes Ly6G-CD11b+ 10.66%
AmCyan-Ly6G
PE-CD11b
D
Neutrophils per cornea
ns
C57BL/6
NLRP3-/-
E
C57BL/6
Neutrophils Ly6G+CD11b+ 91.92%
Monocytes Ly6G-CD11b+ 6.74%
BV510-Ly6G
PE-CD11b
F
gsdmd-/-
Neutrophils Ly6G+CD11b+ 88.42%
Monocytes Ly6G-CD11b+ 9.17%
BV510-Ly6G
PE-CD11b
G
BL/6
gsdmd-/-
Neutrophils per cornea
ns
ns
Mock
24 hr
48 hr
H
IL-1β (pg/cornea)
ns
C57BL/6
gsdmd-/-
I
C57BL/6
Caspase-1/11-/-
Elane-/-
GFP-PAO1
BF
GFP
J
Mean Intensity (gray value/pixel)
ns
***
****
C57BL/6
caspase-1/11-/-
Elane-/-
K
CFU/eye (24 h p.i.)
ns
****
****
****
Inoculum
C57/BL6
caspase-1/11-/-
Elane-/-

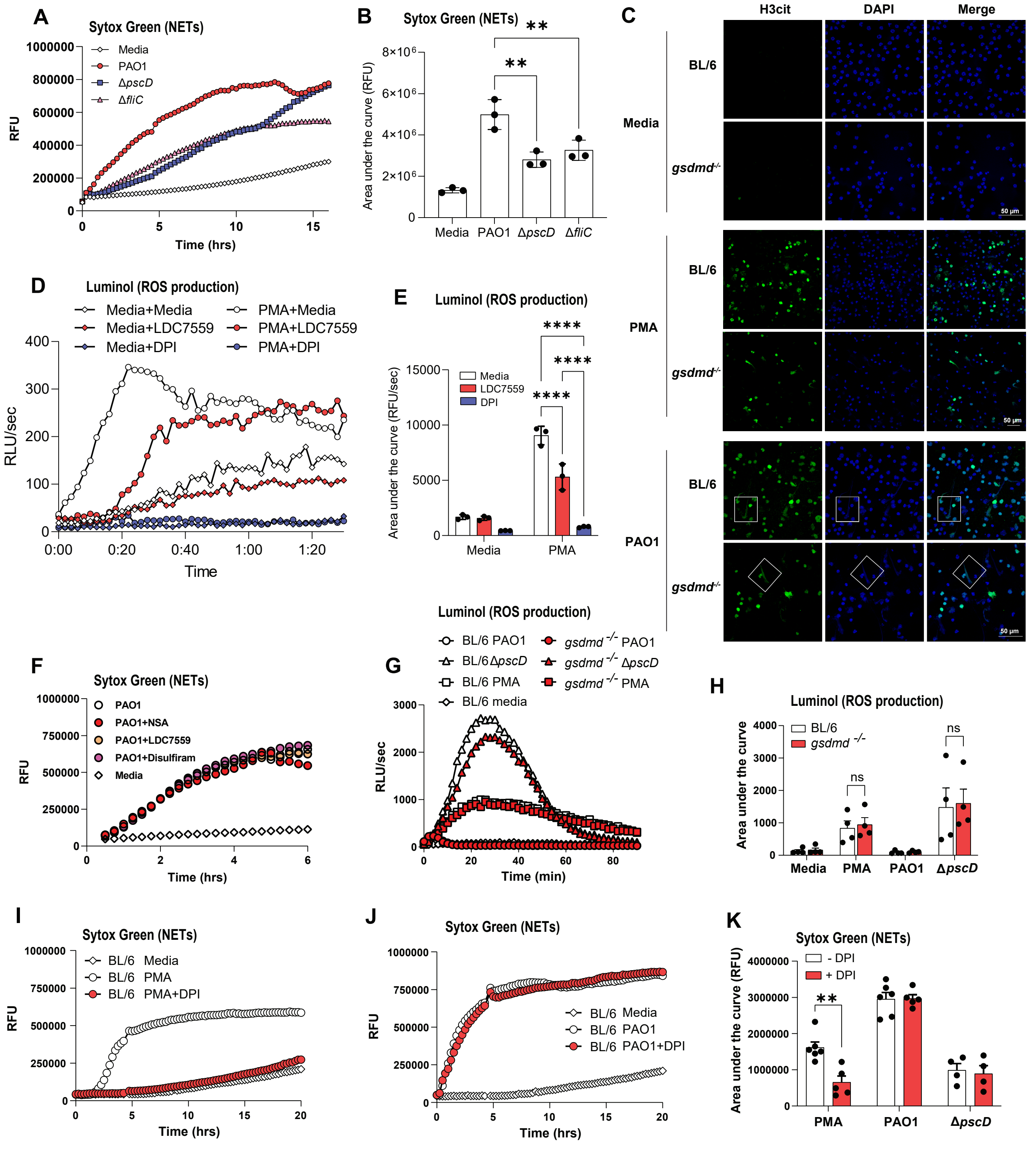
A
Sytox Green (NETs)
Media
PAO1
ΔpscD
ΔfliC
RFU
Time (hrs)
B
Sytox Green (NETs)
Area under the curve (RFU)
Media
PAO1
ΔpscD
ΔfliC
C
H3cit
DAPI
Merge
BL/6
Media
gsdmd-/-
PMA
PAO1
50 μm
D
Luminol (ROS production)
Media+Media
Media+LDC7559
Media+DPI
PMA+Media
PMA+LDC7559
PMA+DPI
RLU/sec
Time
E
Luminol (ROS production)
Media
LDC7559
DPI
Area under the curve (RFU/sec)
F
Sytox Green (NETs)
PAO1
PAO1+NSA
PAO1+LDC7559
PAO1+Disulfiram
Media
RFU
Time (hrs)
G
Luminol (ROS production)
BL/6 PAO1
BL/6 ΔpscD
BL/6 PMA
BL/6 media
gsdmd -/- PAO1
gsdmd -/- ΔpscD
gsdmd -/- PMA
RLU/sec
Time (min)
H
Luminol (ROS production)
BL/6
gsdmd -/-
ns
Area under the curve
Media
PMA
PAO1
ΔpscD
I
Sytox Green (NETs)
BL/6 Media
BL/6 PMA
BL/6 PMA+DPI
RFU
Time (hrs)
J
Sytox Green (NETs)
BL/6 Media
BL/6 PAO1
BL/6 PAO1+DPI
RFU
Time (hrs)
K
Sytox Green (NETs)
- DPI
+ DPI
Area under the curve (RFU)
PMA
PAO1
ΔpscD
